# BAG6-RNF115 Couples Protein Quality Control with Ribosome Assembly

**DOI:** 10.64898/2026.08.06.742952

**Authors:** Anat Raiff, Colin Zenge, Alban Ordureau, Itay Koren

## Abstract

Protein homeostasis relies on protein quality control (PQC) pathways that survey the proteome to eliminate aberrant polypeptides. The BAG6 complex is a central PQC factor that recognizes exposed hydrophobic regions, a feature commonly associated with misfolded, mislocalized, and mistranslated proteins. Whether this surveillance machinery also regulates intact, functional proteins as part of physiological proteostasis has remained unclear. Using unbiased quantitative proteomics, we identify the ribosomal protein RPL22L1 as an endogenous BAG6 substrate whose abundance is controlled by continuous proteasomal degradation. This turnover requires the RNF115 E3 ligase activity but not the canonical BAG6 partner RNF126, defining RPL22L1 as a selective RNF115-dependent substrate. Mechanistically, we map a bipartite hydrophobic degron that distinguishes RPL22L1 from its stable paralog RPL22, and show that BAG6-RNF115-mediated degradation is governed by substrate assembly state. Accordingly, RPL22L1 is protected from degradation upon incorporation into the 60S ribosome, where it substitutes for RPL22. When RPL22 is lost, either genetically or through recurrent inactivating mutations in microsatellite-unstable cancers, the vacant ribosomal binding site permits RPL22L1 incorporation, protecting it from BAG6-mediated degradation. These findings establish unassembly-coupled degradation as a mechanism by which BAG6 regulates the abundance of functional protein components, ensuring that they accumulate only when incorporated into their native macromolecular complexes.

## INTRODUCTION

Protein homeostasis (proteostasis) is crucial for cellular and organismal health, and its disruption underlies a wide range of human diseases, including neurodegeneration, cancer, and immune dysfunction ^1–4^. Proteostasis is maintained by an elaborate protein quality control (PQC) network of molecular chaperones and protein degradation factors that continually monitor protein folding, localization, and abundance throughout the cell^5–7^. To eliminate aberrant or unwanted proteins, PQC machineries collaborate with the ubiquitin-proteasome system (UPS), in which substrates are tagged with polyubiquitin chains by a sequential E1-E2-E3 enzymatic cascade and delivered to the proteasome for degradation^5,6^. To function effectively, these machineries must distinguish their substrates from the broader cellular proteome, a task typically accomplished by E3 ligases that recognize shared determinants within aberrant proteins through short sequence motifs or structural features known as degrons^8–11^. In other cases, chaperones recognize exposed hydrophobic residues on their clients and recruit dedicated E3 ligases to mediate substrate ubiquitination and proteasomal degradation^12,13^.

One of the best-characterized examples of this chaperone-coupled degradation mechanism is the Bcl-2-associated athanogene 6 (BAG6) complex, which serves as a major hub of cytosolic PQC^14,15^. First described as a chaperone that channels nascent tail-anchored proteins to the GET/TRC40 pathway for ER membrane insertion^16^, BAG6 was also shown to route hydrophobic clients to the proteasome by targeting mislocalized membrane and secretory proteins for ubiquitin-dependent degradation^17^, eliminating defective polypeptides released from stalled translation^18^, promoting the degradation of hydrophobic polypeptides translated from noncoding genomic regions^19^ and clearing aberrant readthrough products bearing hydrophobic C-terminal extensions^20^.

A striking feature of the BAG6 substrate landscape is that nearly all known substrates are defective proteins that share a common feature: hydrophobic sequences abnormally exposed to the cytosol^17,18,21,22^. Very few intact proteins have been described as BAG6 substrates. The clearest exceptions are Rab8a and Rab9a, in which BAG6 selectively recognizes the GDP-bound conformation through hydrophobic residues transiently exposed, thereby driving their regulated turnover and regulating vesicular trafficking pathways^23–25^. These examples raise the intriguing possibility that BAG6-dependent degradation can target intact, non-defective endogenous proteins as a mechanism to regulate their steady-state abundance and thereby control diverse cellular pathways. Whether additional physiological endogenous proteins are subject to this type of BAG6-dependent turnover, and what features molecularly distinguish them from the broader cytosolic proteome, remains largely unknown.

To drive substrate degradation, BAG6 harnesses the UPS via its N-terminal ubiquitin-like (UBL) domain, which serves as a platform for recruiting RING-type E3 ligases^17,26^. The best-characterized partner is RNF126, the primary BAG6-dependent E3 ligase that mediates the ubiquitination of aberrant hydrophobic clients^22,26,27^. A structurally related E3 ligase, RNF115, has also been shown to interact with the BAG6 UBL domain^22,26^, however, no substrate has yet been shown to be specifically targeted by BAG6 through RNF115, leaving open the question of whether RNF115 acts redundantly with RNF126 or instead regulates a distinct class of substrates.

To address these two interconnected gaps, whether BAG6 targets endogenous functional proteins beyond defective PQC substrates, and whether RNF115 mediates degradation of a distinct class of BAG6 clients, we used quantitative proteomics to systematically identify novel endogenous substrates of the BAG6 complex. This unbiased approach uncovered RPL22L1, a paralog of the 60S ribosomal protein RPL22^28,29^, as a highly selective BAG6 client. We show that the steady-state abundance of RPL22L1 is regulated by BAG6-mediated proteasomal degradation, and that this regulation operates through the E3 ligase RNF115 and requires its catalytic activity, establishing RPL22L1 as the first dedicated, RNF115-specific substrate of the BAG6 pathway. We found that BAG6-mediated degradation of RPL22L1 operates in both the cytosol and the nucleus, ensuring dual-compartment control of its abundance. In addition, we mapped a bipartite degron that distinguishes RPL22L1 from its stable paralog RPL22, and show that its hydrophobic residues are essential for recognition, conforming to the hydrophobic degron motif characteristic of BAG6 clients. Finally, we demonstrate that incorporation into the 60S ribosome shields RPL22L1 from BAG6 recognition, coupling its turnover to its functional state. Upon loss of RPL22, RPL22L1 assembles into the ribosome and escapes degradation, and the same mechanism operates in cancer cells carrying inactivating RPL22 mutations, establishing a rheostat-like mechanism in which RPL22 availability tunes the balance between RPL22L1 degradation and preservation. Together, these findings expand the BAG6 substrate repertoire beyond defective proteins, assign a distinct cellular function to the BAG6-RNF115 axis, and reveal a PQC mechanism in which a ribosomal protein is continuously degraded unless engaged in its physiological complex.

## RESULTS

### RPL22L1 is a novel BAG6 substrate identified by quantitative proteomics

To identify novel endogenous substrates of the BAG6 pathway, we performed tandem mass tag (TMT)-based quantitative proteomics to compare wild-type (WT) and BAG6-deficient human embryonic kidney 293T (HEK293T) cells generated using CRISPR-Cas9. To ensure the reliability of our findings and minimize potential off-target effects of CRISPR-Cas9 editing, we generated two independent BAG6-deficient cell lines using distinct single guide RNAs (sgRNAs). Proteomics analysis across all three conditions (WT, BAG6 knockout (KO)1, and BAG6 KO2) identified a total of 9,940 proteins **(Fig. 1a and Supplementary Table 1)**. Principal component analysis (PCA) confirmed clear separation between WT and BAG6 KO samples **(Supplementary Fig. 1a)**. Statistical analysis using Welch’s t-test, with significance defined as a Benjamini-Hochberg (BH) false discovery rate (FDR) below 5% combined with a fold change greater than ±2, identified 51 proteins whose abundance was significantly altered following BAG6 loss, with 31 increased and 20 decreased in abundance **(Fig. 1a)**. Among these, RPL22L1 was the most significantly increased protein following BAG6 depletion, and was consistently identified in both independent KO lines **(Fig. 1a-b)**.

**Fig. 1.**
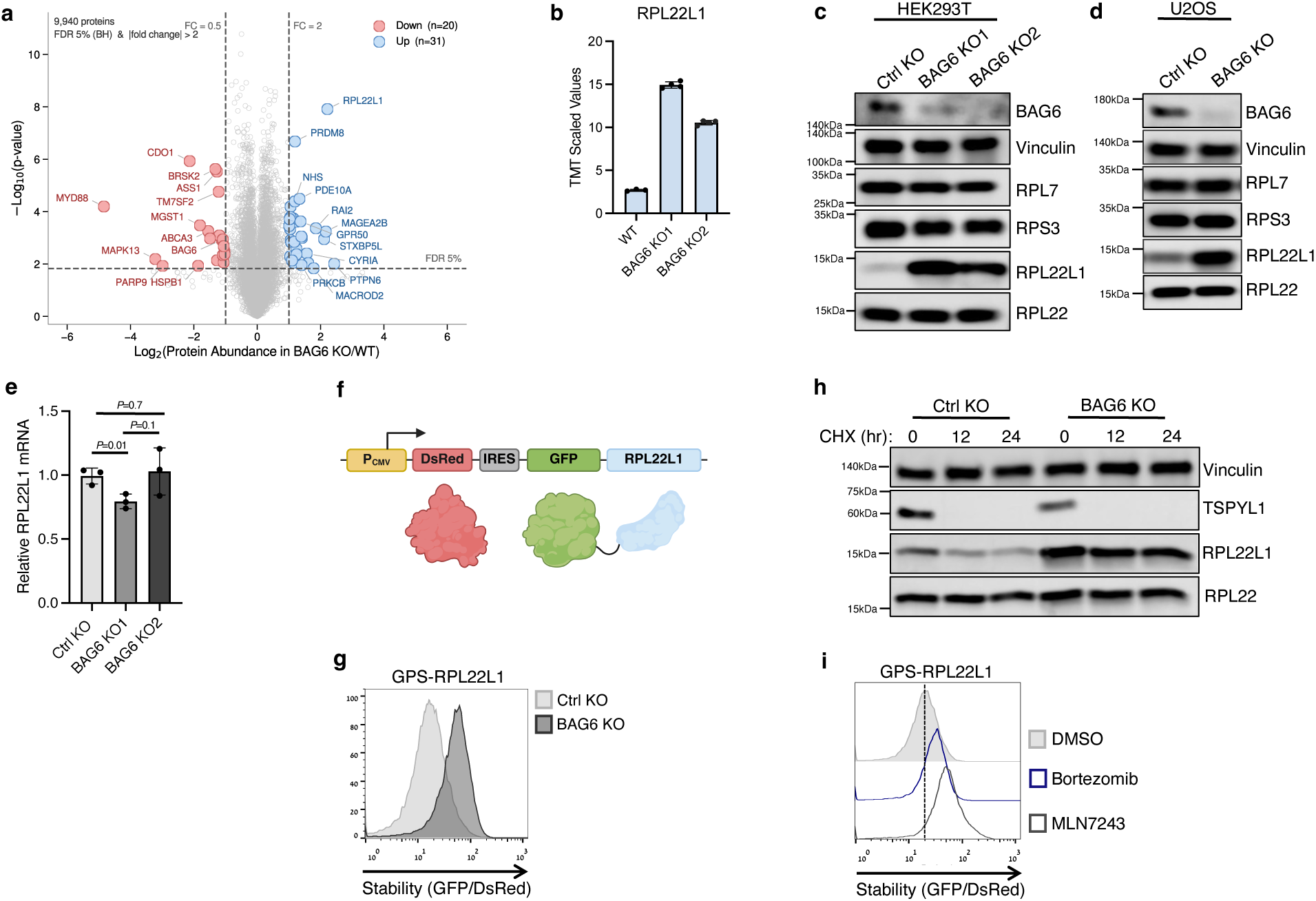
BAG6 mediates proteasomal degradation of RPL22L1. **a,** Volcano plot of log₂ fold change versus -log₁₀(*P* value) from TMT-based proteomic analysis of the total proteome, comparing control parental (WT) HEK293T cells with two independent BAG6 KO lines. *P* values were calculated by Welch’s *t*-test and corrected using the Benjamini–Hochberg procedure; proteins with FDR < 5% and |fold change| > 2 were considered significantly changed, yielding 31 increased (light blue) and 20 decreased (light red) proteins out of 9,940 quantified. Selected proteins are labeled. RPL22L1 is the most significantly increased protein upon BAG6 loss; the reduction in BAG6 itself confirms its depletion. **b,** TMT relative abundance (scaled values) of RPL22L1 in WT and the two independent BAG6 KO lines. Data are mean ± s.e.; *n* = 3 (WT) and *n* = 4 (each BAG6 KO) biological replicates. Quantification based on 11 peptides. **c,** Immunoblot analysis of BAG6, RPL7, RPS3, RPL22 and RPL22L1 in control (Ctrl) KO and two independent BAG6 KO HEK293T lines. Ctrl KO, sgRNA targeting AAVS1. Vinculin served as a loading control. **d,** Immunoblot analysis of BAG6, RPL7, RPS3, RPL22 and RPL22L1 in Ctrl KO and BAG6 KO U2OS cells. Vinculin served as a loading control. **e,** Quantitative RT-PCR of RPL22L1 mRNA in Ctrl KO and two independent BAG6 KO cells. Data are mean ± s.e. of *n* = 3 for Ctrl KO and *n* = 3 for both BAG6 KO1 and KO2, technical replicates. *P* value determined by two-tailed unpaired *t*-test. **f,** Schematic of the bicistronic global protein stability (GPS) lentiviral reporter, in which DsRed and GFP-RPL22L1 are expressed from a single transcript, with GFP-RPL22L1 translated via an IRES. The GFP/DsRed ratio measured by flow cytometry reports RPL22L1 stability. **g,** Flow cytometry analysis of the GPS-RPL22L1 reporter in Ctrl KO and BAG6 KO1 cells. A rightward shift indicates increased GFP-RPL22L1 stability. **h,** Ctrl KO and BAG6 KO1 cells were treated with 50 μg ml⁻¹ cycloheximide (CHX) for the indicated times, followed by immunoblotting for RPL22 and RPL22L1. Vinculin served as a loading control. TSPYL1, a short-lived protein, served as a positive control for effective translational inhibition; RPL22 served as a stable-paralog control. hr: hours. **i,** Flow cytometry analysis of the GPS-RPL22L1 reporter in cells treated with vehicle control (DMSO), 1 µM bortezomib or 1 µM MLN7243 for 6 hr.

RPL22L1 is the paralog of the ribosomal protein RPL22, and both are components of the 60S large ribosomal subunit^28,29^. Strikingly, RPL22L1 was the only one significantly increased upon BAG6 loss among the 90 ribosomal proteins detected in the proteomics analysis (41 of the 40S and 49 of the 60S subunit) **(Supplementary Fig. 1b)**. Examination of individual 60S and 40S ribosomal proteins from the proteomics analysis confirmed that levels of other ribosomal proteins, including the paralog RPL22, were not increased upon BAG6 depletion **(Supplementary Fig. 1c)**, highlighting the selective regulation of RPL22L1 by BAG6 across the ribosomal protein repertoire.

### BAG6 regulates RPL22L1 turnover via the UPS

To validate the proteomics findings at the endogenous level, we examined RPL22L1 protein levels by western blot. In HEK293T cells, both independent BAG6 KO lines (KO1 and KO2) showed efficient loss of BAG6 protein accompanied by a marked accumulation of RPL22L1 relative to control (Ctrl) KO cells, validating the proteomics analysis **(Fig. 1c)**. By contrast, other ribosomal proteins of both the large 60S subunit (RPL7, RPL22) and the small 40S subunit (RPS3) remained unchanged **(Fig. 1c)** indicating that BAG6-mediated regulation is specific to RPL22L1 rather than reflecting a general effect on ribosomal proteins. To exclude the possibility of a cell-type-specific phenotype, we extended this analysis to U2OS cells, in which efficient BAG6 depletion resulted in RPL22L1 accumulation **(Fig. 1d)**. Notably, quantitative real-time (RT)-PCR revealed that RPL22L1 mRNA levels were unchanged in two independent BAG6 KO cells **(Fig. 1e)**, suggesting that BAG6 regulates RPL22L1 at the post-translational level, likely by promoting its degradation.

To investigate whether RPL22L1 undergoes BAG6-dependent degradation, we employed the global protein stability (GPS) dual-fluorescence reporter system^10,11,30^. To this end, RPL22L1 was fused to GFP and co-expressed with an internal DsRed control from the same transcript, allowing quantitative measurement of relative protein stability by flow cytometry based on the GFP/DsRed fluorescence ratio **(Fig. 1f)**. Using this assay, we found that GPS-RPL22L1 was substantially stabilized in BAG6 KO cells compared to control cells **(Fig. 1g)**, consistent with the endogenous protein data **(Fig. 1c-d)**. To rule out the possibility that the GFP tag non-specifically renders ribosomal proteins susceptible to BAG6-mediated degradation, we generated GPS reporters for additional ribosomal proteins, including RPL7 and RPL22. Neither GPS-RPL7 nor GPS-RPL22 was stabilized in BAG6 KO cells **(Supplementary Fig. 1d)**. Furthermore, GFP tagging did not impair ribosome incorporation **(Supplementary Fig. 1e),** indicating that the observed BAG6-dependent degradation of GPS-RPL22L1 is not an artifact of the fusion construct. In line with these findings, cycloheximide (CHX) chase assays demonstrated that RPL22L1 turnover is regulated by BAG6, where RPL22L1 levels decreased upon CHX treatment in control KO cells, but its degradation was impaired in BAG6-deficient cells **(Fig. 1h)**. By contrast, its paralog RPL22 was stable under CHX treatment in both backgrounds, confirming that BAG6-dependent turnover is specific to RPL22L1 **(Fig. 1h)**. TSPYL1, a known short-lived protein^10^, served as a positive control to confirm effective translational inhibition and was rapidly degraded in both KO backgrounds **(Fig. 1h)**. Consistent with our data, reanalysis of a degradomics dataset^31^ identified RPL22L1 as the most rapidly degraded among all 75 ribosomal proteins quantified in this study **(Supplementary Fig. 1f)**, independently corroborating that RPL22L1 undergoes exceptionally high turnover compared to the human ribosomal protein repertoire. Finally, we found that RPL22L1 degradation proceeds through the UPS. Treatment of the GPS-RPL22L1 reporter cells with the E1 ubiquitin-activating enzyme inhibitor MLN7243 or the proteasome inhibitor bortezomib led to marked stabilization of GPS-RPL22L1 **(Fig. 1i)**, demonstrating that RPL22L1 turnover requires both ubiquitination and proteasomal activity. Notably, the degree of stabilization observed in BAG6 KO cells was comparable to that achieved by treatment with bortezomib **(Supplementary Fig. 1g)**, indicating that BAG6 is the primary, and likely sole, route through which RPL22L1 is delivered to the proteasome for degradation.

Together, these findings demonstrate that BAG6 selectively targets RPL22L1 for proteasome- and ubiquitin-dependent degradation, thereby controlling its turnover.

### The BAG6-RNF115 axis mediates proteasomal degradation of RPL22L1

We next investigated whether RPL22L1 physically interacts with the BAG6 complex. Co-immunoprecipitation (co-IP) of HA-tagged BAG6 recovered endogenous RPL22L1 but not RPL22, indicating that BAG6 selectively associates with RPL22L1 **(Fig. 2a)**. To further dissect this interaction, we examined the role of the UBL domain of BAG6, which is known to mediate the recruitment of the E3 ligases RNF126 and RNF115 to the BAG6 complex^22^. For this purpose, we generated a BAG6 mutant lacking the UBL domain (ΔUBL) and performed co-IP experiments, revealing that RPL22L1 binding was retained in the ΔUBL mutant at levels comparable to WT BAG6 **(Fig. 2a)**. This indicates that BAG6 recognizes RPL22L1 through a region distinct from the UBL domain, consistent with the UBL domain functioning as an E3 ligase recruitment module rather than a substrate-binding domain. In contrast, interactions with RNF115 and RNF126 were abolished in the ΔUBL mutant **(Fig. 2a)**, confirming that the UBL domain specifically mediates E3 ligase engagement.

**Fig. 2.**
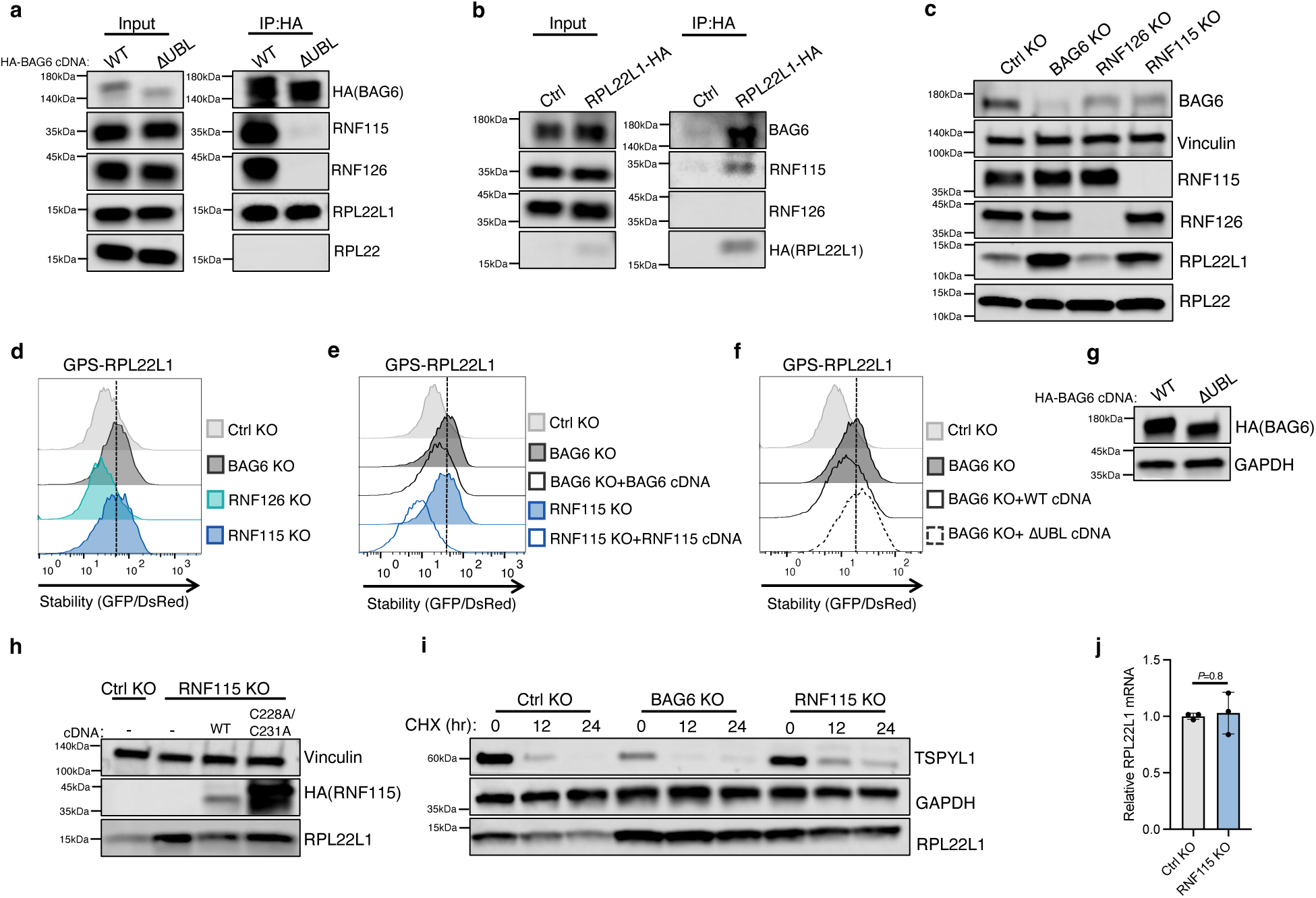
The BAG6-RNF115 complex regulates RPL22L1 degradation. **a,** HEK293T cells stably expressing HA-tagged BAG6 (WT or ΔUBL, lacking the N-terminal ubiquitin-like domain) were pre-treated with 1 µM bortezomib for 6 hr, followed by HA immunoprecipitation (IP:HA). Interactions with RNF115, RNF126, RPL22L1 and RPL22 were assessed by immunoblotting of immunoprecipitates and total cell extracts (Input). **b,** HEK293T cells expressing HA-tagged RPL22L1 were subjected to HA immunoprecipitation. Association with endogenous BAG6, RNF126, and RNF115 was assessed by immunoblotting of immunoprecipitates (IP:HA) and total cell extracts (Input). **c,** Immunoblot analysis of BAG6, RNF115, RNF126, RPL22 and RPL22L1 in Ctrl KO, BAG6 KO1, RNF126 KO and RNF115 KO cells. Vinculin served as a loading control. **d,** GPS assay of the GPS-RPL22L1 reporter in different KO backgrounds as assessed by flow cytometry. **e,** Flow cytometry analysis of the GPS-RPL22L1 reporter stability in Ctrl KO, BAG6 KO1 and RNF115 KO cells, and in the corresponding KO cells reconstituted with WT BAG6 or WT RNF115 cDNA. **f,g,** Flow cytometry analysis of the GPS-RPL22L1 reporter in Ctrl KO and BAG6 KO1 cells reconstituted with WT BAG6 or the ΔUBL mutant **(f)**, with expression of the exogenous HA-tagged BAG6 constructs confirmed by immunoblotting **(g)**. GAPDH served as a loading control. **h,** Immunoblot of Ctrl KO and RNF115 KO cells reconstituted with WT RNF115 or the RING-domain mutant C228A/C231A, probed with antibodies against HA (HA-RNF115), and RPL22L1. Vinculin served as a loading control. **i,** CHX chase assay of RPL22L1 in different KO backgrounds. Cells were treated with 50 μg ml⁻¹ CHX for the indicated times, followed by immunoblotting for TSPYL1 and RPL22L1. TSPYL1 is a short-lived protein that served as a control for efficient translation inhibition. GAPDH served as a loading control. hr: hours. **j,** Quantitative RT-PCR of RPL22L1 mRNA in Ctrl KO and RNF115 KO cells. Data are mean ± s.e. of *n* = 3 technical replicates, representative of three independent experiments. *P* value determined by two-tailed unpaired *t*-test.

BAG6 is known to function with two E3 ligases: the canonical partner RNF126 and the more recently characterized RNF115. RNF115 and RNF126 are homologous ligases that share the same domain architecture, including a conserved N-terminal zinc-finger (ZNF) domain and a C-terminal RING domain **(Supplementary Fig. 2a-b)**, and both associate with BAG6, with RNF126 exhibiting greater BAG6 co-immunoprecipitation than RNF115 **(Supplementary Fig. 2c)**. To determine which E3 ligase mediates RPL22L1 degradation, we immunoprecipitated HA-tagged RPL22L1 and probed for BAG6, RNF115, and RNF126. Surprisingly, despite its weaker interaction with BAG6 **(Supplementary Fig. 2c)**, RNF115 associated with RPL22L1, whereas RNF126 did not **(Fig. 2b)**.

These findings are intriguing, as to date no specific substrate for the BAG6-RNF115 pathway has been described, and to our knowledge RPL22L1 is the first RNF115-specific substrate, which is not redundantly targeted by RNF126. Having established that RPL22L1 interacts with BAG6 and RNF115, we next sought to functionally delineate the degradation pathway responsible for RPL22L1 turnover. To this end, we generated RNF115- and RNF126-deficient cells using CRISPR-Cas9. Western blot analysis of endogenous RPL22L1 in BAG6, RNF115, and RNF126 KO cells revealed that RPL22L1 accumulates specifically in BAG6 and RNF115 KO cells, whereas RNF126 KO had no effect **(Fig. 2c)**. These results were supported by the GPS reporter system, in which GPS-RPL22L1 was stabilized in BAG6- or RNF115-deficient cells but not upon RNF126 loss **(Fig. 2d)**. Rescue experiments further confirmed pathway specificity, as re-introduction of WT BAG6 cDNA into BAG6 KO cells, or WT RNF115 cDNA into RNF115 KO cells, restored RPL22L1 degradation as measured by the GPS assay **(Fig. 2e)**. Importantly, re-expression of the BAG6 ΔUBL mutant failed to rescue RPL22L1 degradation in BAG6 KO cells **(Fig. 2f)**, despite being expressed at levels comparable to WT BAG6 **(Fig. 2g),** demonstrating that while the UBL domain is dispensable for RPL22L1 binding **(Fig. 2a)**, it is essential for its degradation. These findings support a model in which BAG6 recognizes RPL22L1 independently of the UBL domain but requires the UBL domain to recruit the E3 ligase RNF115, which in turn ubiquitinates RPL22L1 and targets it for proteasomal degradation. Supporting this model, we found that the ubiquitin ligase activity of RNF115 is essential for RPL22L1 degradation. Whereas re-expression of WT RNF115 cDNA in RNF115 KO cells restored RPL22L1 protein levels to those of control cells, the RING mutant C228A/C231A failed to do so despite being expressed at substantially higher levels than WT **(Fig. 2h)**.

To assess the effect of the E3 ligase RNF115 on RPL22L1 protein turnover, we performed CHX chase assays. In control KO cells, RPL22L1 displayed progressive degradation over the time course, whereas its turnover was markedly attenuated in both BAG6- and RNF115-deficient cells **(Fig. 2i)**. These results demonstrate that the BAG6-RNF115 pathway actively contributes to RPL22L1 protein turnover. To rule out the possibility of transcriptional regulation of RPL22L1 by RNF115, we performed quantitative RT-PCR. RPL22L1 mRNA levels were unchanged in RNF115 KO cells **(Fig. 2j)**, confirming that BAG6-RNF115 regulates RPL22L1 exclusively through post-translational protein degradation.

### The BAG6 pathway targets both nuclear and cytoplasmic RPL22L1 for degradation

BAG6 harbors a functional C-terminal nuclear localization signal (NLS)^32^ and is known to be distributed between the nucleus and the cytoplasm^33,34^. RPL22L1 has also been reported to be present in both compartment^29^. Having established that BAG6 and RNF115 mediate RPL22L1 degradation, we therefore asked in which cellular compartment this regulation occurs. Fractionation of cells into cytosolic and nuclear fractions showed that BAG6, RNF115 and RNF126 were present in both compartments, with relative enrichment in the cytosol, while RPL22L1 and RPL22 were present at comparable levels in the cytosol and nucleus **(Fig. 3a)**. Thus, all components of the pathway are available in both cellular compartments, consistent with the possibility that RPL22L1 degradation occurs in both the cytoplasm and the nucleus.

**Fig. 3.**
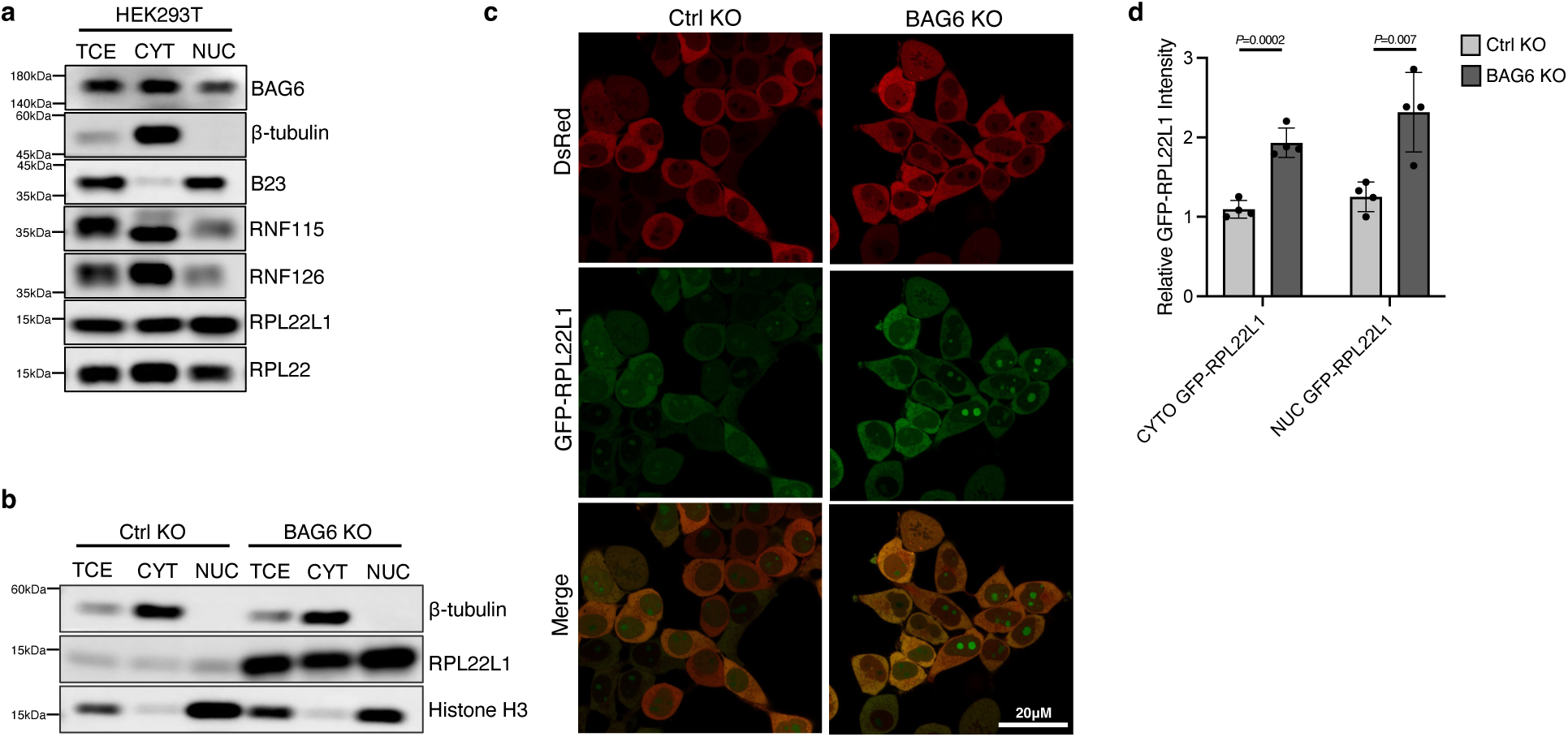
Both nuclear and cytoplasmic RPL22L1 are targeted for degradation by the BAG6 pathway. **a,** Immunoblot analysis of BAG6, RNF115, RNF126, RPL22 and RPL22L1 in total cell extract (TCE), cytosolic (CYT) and nuclear (NUC) fractions of HEK293T cells. To confirm fractionation purity, β-tubulin served as a cytosolic marker and B23 as a nuclear marker. **b,** Immunoblot analysis of RPL22L1 in TCE, CYT and NUC fractions of Ctrl KO and BAG6 KO cells. β-tubulin and histone H3 served as cytoplasmic and nuclear fractionation controls, respectively. **c,** Representative confocal microscopy images of Ctrl KO and BAG6 KO1 cells expressing the GPS-RPL22L1 reporter. GFP–RPL22L1 (green), DsRed (red), and the merged image are shown. Scale bar, 20 µm. **d,** Quantification of GFP-RPL22L1 fluorescence intensity in the cytoplasmic (CYTO) and nuclear (NUC) compartments of Ctrl KO and BAG6 KO1 cells, normalized to DsRed. DsRed, which fills the cytoplasm but is excluded from the nucleus, was used both to delineate the two compartments and as an internal reference for reporter expression. Each data point represents the mean of 10–15 cells within one field; data are mean ± s.d. of *n* = 4 fields. *P* values determined by two-tailed unpaired *t*-test.

We next asked where RPL22L1 is stabilized upon BAG6 loss. Fractionation of control and BAG6 KO cells revealed accumulation of RPL22L1 in both the cytosolic and nuclear fractions **(Fig. 3b)**. These findings were corroborated by confocal imaging analysis of GPS-RPL22L1 reporter cells, which showed increased GFP-RPL22L1 fluorescence intensity in BAG6 KO compared to control KO cells, with quantification confirming significant stabilization in both compartments **(Fig. 3c-d)**.

Together, these results indicate that BAG6 recognizes RPL22L1 in both the cytoplasm and the nucleus, pointing to tight surveillance of RPL22L1 across both compartments.

### Identification of the RPL22L1 degron recognized by BAG6

RPL22L1 and RPL22 share 70% amino acid sequence identity, yet only RPL22L1 is targeted for BAG6-mediated degradation. Pairwise alignment of the two proteins showed extensive sequence conservation throughout much of the coding region, with divergence confined to several discrete stretches **(Supplementary Fig. 3a)**. Structural prediction using AlphaFold 3^35^ indicated that RPL22L1 and RPL22 adopt highly similar three-dimensional folds **(Supplementary Fig. 3b)**, suggesting that substrate selectivity is dictated by sequence differences rather than global structural features. We therefore sought to identify the RPL22L1 degron recognized by BAG6.

To map the degron, we generated chimeric constructs in which discrete segments of RPL22L1 were replaced by the corresponding regions of RPL22. RPL22L1 was divided into an N-terminal (aa 1-33), a middle (aa 39-71) and a C-terminal (aa 99-122) segment, each of which was individually swapped with the corresponding regions of RPL22 while the intervening linkers were left unchanged, yielding the chimeras RPL22L1 (N)22, RPL22L1 (M)22 and RPL22L1 (C)22 **(Fig. 4a and Supplementary Fig. 3c-e)**. Using the GPS reporter system, we assessed the stability of each chimera in control and BAG6 KO cells by flow cytometry. Replacement of either the N-terminal or the middle region abolished BAG6-dependent degradation, whereas RPL22L1 (C)22 continued to be degraded in a BAG6-dependent manner **(Fig. 4b)**. Thus, the degron resides within the N-terminal and middle regions, precisely the segments that diverge most from RPL22 **(Supplementary Fig. 3a),** and both are required for BAG6-dependent degradation in the context of the full-length protein.

**Fig. 4.**
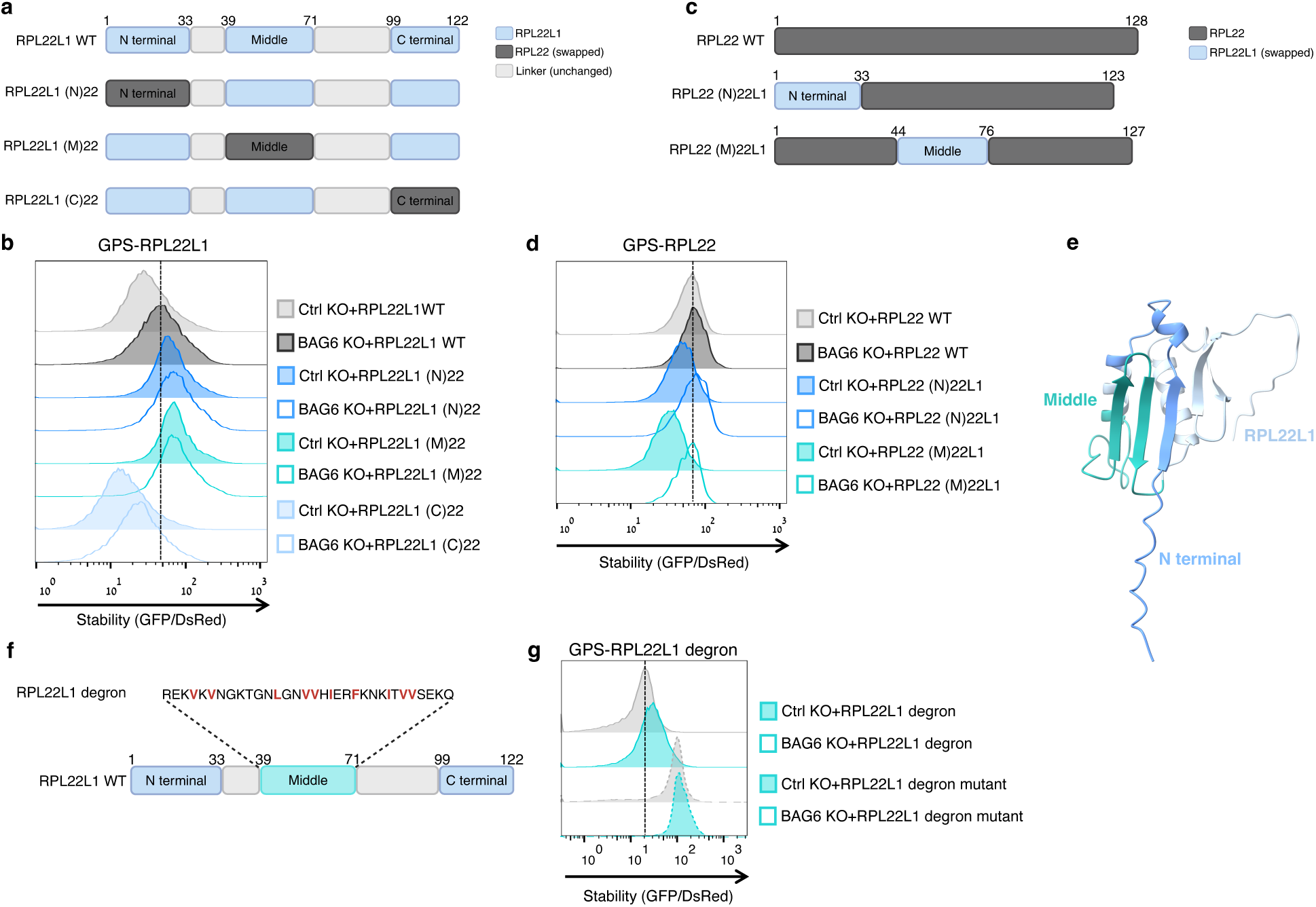
A bipartite degron in RPL22L1 is required for BAG6-mediated degradation. **a,** Schematic illustration of the RPL22L1 chimeras. The N-terminal (aa 1-33), middle (aa 39-71) or C-terminal (aa 99-122) region of RPL22L1 was replaced with the corresponding RPL22 sequence, generating RPL22L1(N)22, RPL22L1(M)22 and RPL22L1(C)22. Light blue, RPL22L1; dark grey, RPL22 (swapped); light grey, unchanged linker regions. **b,** Stability analysis of GPS-fused full-length RPL22L1 (WT) or the (N)22, (M)22 and (C)22 chimeras in Ctrl KO and BAG6 KO1 cells. **c,** Schematic illustration of the reciprocal chimeras, in which the RPL22L1 N-terminal or middle region was transplanted into the corresponding position of RPL22, generating RPL22(N)22L1 and RPL22(M)22L1. Dark grey, RPL22; light blue, RPL22L1 (swapped). **d,** Flow cytometry analysis of GPS-fused full-length RPL22 (WT) or the (N)22L1 and (M)22L1 chimeras in Ctrl KO and BAG6 KO1 cells. **e,** AlphaFold-predicted structural model of RPL22L1 highlighting the N-terminal region (aa 1-33, blue) and the middle region (aa 39-71, teal) identified as required for BAG6-mediated degradation. Visualized in ChimeraX. **f,** Illustration of the RPL22L1 middle degron (aa 39-71) sequence (RPL22L1 degron), with hydrophobic residues highlighted in red. **g,** Stability analysis of GPS reporters fused to the WT RPL22L1 middle degron or to the degron mutant in which the hydrophobic residues were substituted, in Ctrl KO and BAG6 KO1 cells.

To exclude the possibility that the stabilization of the RPL22L1 (N)22 and (M)22 chimeras resulted from altered localization rather than loss of BAG6 recognition, we examined the localization of each variant by immunofluorescence using the nucleolar marker B23 and the cytoplasmic marker DsRed. WT RPL22L1 and all three chimeras co-localized with both markers **(Supplementary Fig. 4a)**, indicating that none of the sequence swaps disrupted proper subcellular targeting.

We next asked whether the N-terminal and middle regions are sufficient to confer BAG6-dependent degradation. To this end, we carried out the reciprocal experiment, transplanting the RPL22L1 N-terminal or middle region into the corresponding position of the stable paralog RPL22 **(Fig. 4c)**. As expected, GPS-RPL22 was insensitive to BAG6 loss **(Fig. 4d)**. In contrast, introduction of either RPL22L1 region was sufficient to render RPL22 susceptible to BAG6-dependent degradation, demonstrating that each region can impose BAG6-mediated turnover on an otherwise stable protein **(Fig. 4d)**. Notably, the middle region produced a stronger effect than the N-terminal region **(Fig. 4d)**, suggesting that although both elements are transferable, the middle region constitutes the more potent degron.

Mapping the two regions onto the AlphaFold 3-predicted structure of RPL22L1 showed that the N-terminal and middle segments form spatially adjacent surfaces on the folded protein **(Fig. 4e)**, consistent with their joint requirement within the intact protein.

We next asked whether either region can act as an autonomous degron outside the native protein. The isolated N-terminal region (aa 1-33), the middle region (aa 39-71), or both together (aa 1-71) were fused to GFP within the GPS reporter **(Supplementary Fig. 3f-h and Supplementary Fig. 4b)**. All three constructs were stabilized in BAG6 KO relative to control KO cells **(Supplementary Fig. 4c)**, demonstrating that each region encodes a transferable degron sufficient to direct BAG6-mediated degradation. Taken altogether, although both regions can act as degrons in isolation, both are required for efficient degradation of the intact protein.

Finally, we sought to define the critical residues within the degron motif. We focused on the middle region because it conferred the strongest BAG6 dependence **(Fig. 4d).** This 33-amino acid segment (aa 39-71) contains several hydrophobic residues distributed along its length **(Fig. 4f).** BAG6 is well established as a sensor of exposed hydrophobicity, and previous proteome-wide internal degron mapping^36^ has defined a BAG6 degron motif in which a stretch of hydrophobic residues are required for degradation, as substitution of these residues stabilizes the corresponding degron models. We therefore reasoned that these hydrophobic residues constitute the BAG6 recognition determinant within RPL22L1. Consistent with this model, mutation of these residues substantially stabilized the GFP-fused mutant degron and abolished its BAG6-dependent degradation, whereas the WT degron remained BAG6-sensitive **(Fig. 4g)**. These hydrophobic positions therefore represent the critical determinants of recognition, and the RPL22L1 degron conforms to the hydrophobic BAG6 degron motif previously defined by proteome-wide screening^36^.

Together, these results define a bipartite degron in RPL22L1, composed of two spatially adjacent regions whose hydrophobic residues serve as the core recognition determinant. The presence of these sequences in RPL22L1 but not in RPL22 provides a molecular explanation for the selective degradation of RPL22L1 by the BAG6 pathway.

### Ribosome incorporation protects RPL22L1 from BAG6-mediated degradation

RPL22 and RPL22L1 functionally compensate for one another and compete for the same binding site on the 60S ribosomal subunit^28^. Having established that BAG6 selectively targets RPL22L1 for degradation, we next asked under what conditions RPL22L1 escapes this surveillance. Because only unassembled ribosomal proteins are expected to be accessible to cytosolic quality control factors, we hypothesized that incorporation of RPL22L1 into mature ribosomes would protect it from BAG6-mediated degradation. Since RPL22 is the predominant paralog across most cell types **(Supplementary Fig. 5a)**, competition from RPL22 is expected to limit RPL22L1 assembly; much of the RPL22L1 pool remains unincorporated and is therefore accessible to BAG6-mediated degradation. This model predicts that depletion of RPL22 should vacate the shared binding site, allowing RPL22L1 to assemble into the 60S subunit and thereby protect it from BAG6-mediated degradation.

To test this model, we generated RPL22 KO cells using two independent sgRNAs. Western blotting confirmed loss of RPL22 protein and revealed a marked increase in endogenous RPL22L1 protein in both HEK293T and HeLa cells **(Fig. 5a)**, whereas the abundance of other ribosomal proteins, including RPL7 and RPS3, remained unchanged **(Supplementary Fig. 5b),** indicating that the effect is specific to RPL22L1 rather than a general consequence of perturbed ribosome composition.

**Fig. 5.**
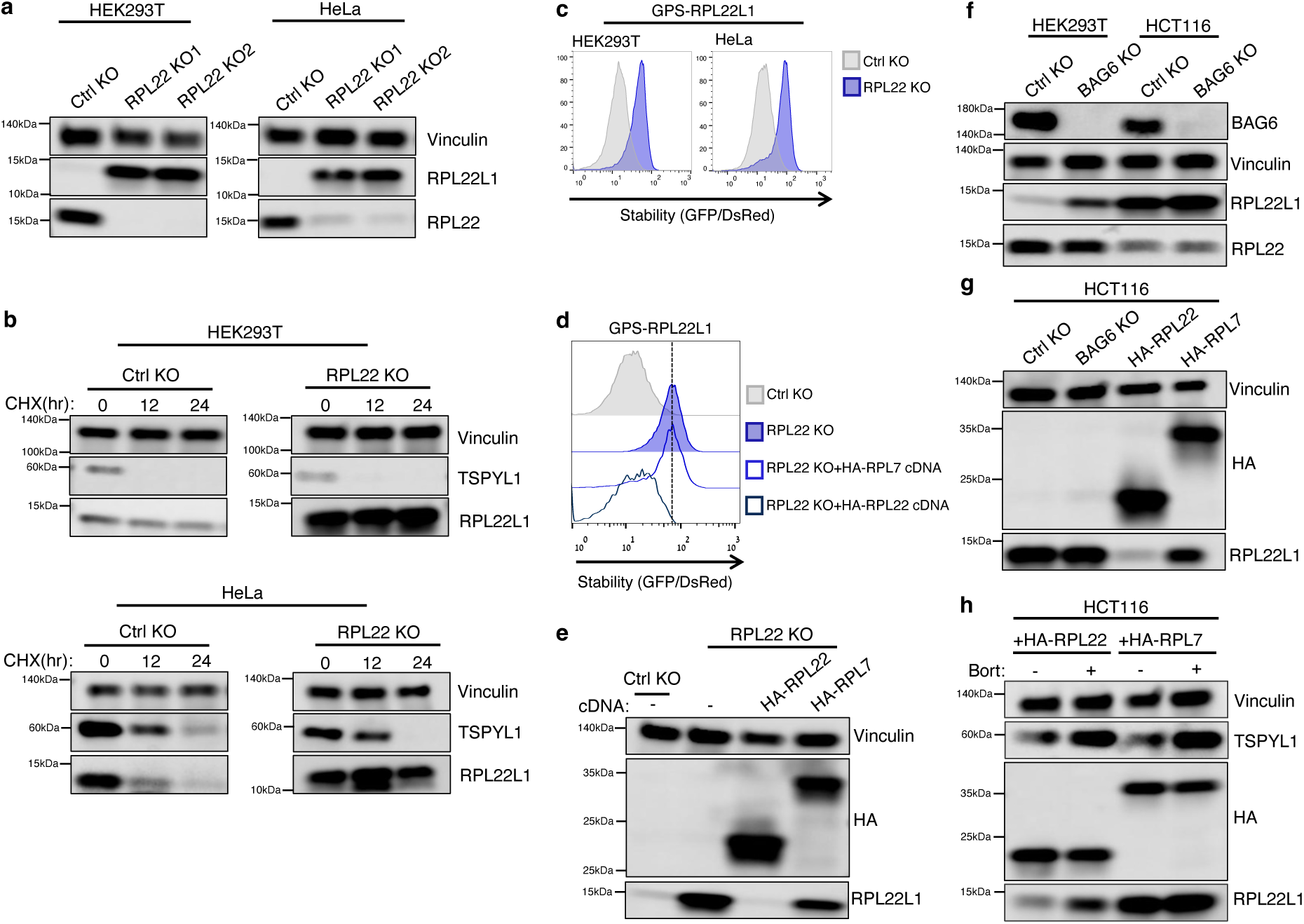
Ribosome incorporation protects RPL22L1 from BAG6-mediated degradation. **a,** Immunoblot analysis of RPL22 and RPL22L1 in Ctrl KO and two independent RPL22 KO HEK293T (left) and HeLa (right) cells. Vinculin served as a loading control. **b,** Ctrl KO and RPL22 KO HEK293T (top) and HeLa (bottom) cells were treated with 50 μg ml⁻¹ CHX for the indicated times, followed by immunoblotting for TSPYL1 and RPL22L1. Vinculin served as a loading control. hr: hours. **c,** Flow cytometry analysis of the GPS-RPL22L1 reporter in Ctrl KO and RPL22 KO HEK293T and HeLa cells. **d,** Flow cytometry analysis of the GPS-RPL22L1 reporter in Ctrl KO cells, RPL22 KO cells, and RPL22 KO cells reconstituted with HA-RPL7 or HA-RPL22 cDNA. **e,** Immunoblot of Ctrl KO and RPL22 KO cells reconstituted with HA-RPL22 or HA-RPL7 cDNA, probed with anti-HA and anti-RPL22L1. Vinculin served as a loading control. **f,** Comparative immunoblot analysis of BAG6, RPL22 and RPL22L1 in Ctrl KO and BAG6 KO1 HEK293T and HCT116 cells. HCT116 cells carry a heterozygous RPL22 frameshift allele (p.K15Rfs*5). Vinculin served as a loading control. **g,** RPL22L1 levels assessed by immunoblotting in HCT116 Ctrl KO cells, BAG6 KO cells, and cells expressing HA-RPL22 or HA-RPL7. Expression of the HA-tagged constructs was verified with an anti-HA antibody. Vinculin served as a loading control. **h,** HCT116 cells reconstituted with HA-RPL22 or HA-RPL7 were treated with DMSO or 1 µM bortezomib (Bort.) for 6 hr, followed by immunoblotting for TSPYL1, HA and RPL22L1. TSPYL1 served as a positive control for effective proteasome inhibition; vinculin served as a loading control.

RPL22 is known to repress RPL22L1 expression at the transcript level^28,37^, which we confirmed in our cells. However, RPL22L1 mRNA levels were only modestly increased in RPL22 KO compared with control KO cells **(Supplementary Fig. 5c)**, suggesting that transcriptional regulation alone may not fully account for the effects of RPL22 loss. We therefore asked whether RPL22 also regulates RPL22L1 at the protein level. CHX chase experiments showed that RPL22L1 protein turnover was markedly attenuated in the absence of RPL22 compared to control KO cells **(Fig. 5b)**. Consistent with this, the GPS reporter that directly measures protein stability revealed marked stabilization of GPS-RPL22L1 in RPL22 KO cells **(Fig. 5c)**. Thus, RPL22 regulates RPL22L1 at both the transcript and protein levels, with regulation of protein stability representing the more prominent mechanism controlling RPL22L1 abundance.

To test whether this stabilization reflects competition for the shared ribosomal site rather than a general requirement for a 60S component, we performed rescue experiments. Re-expression of HA-RPL22 in RPL22 KO cells restored degradation of GPS-RPL22L1 **(Fig. 5d)** and reduced endogenous RPL22L1 levels **(Fig. 5e)**. In contrast, HA-RPL7, a non-paralogous ribosomal protein that occupies a distinct site on the 60S subunit and was expressed at comparable levels, neither restored reporter degradation **(Fig. 5d)** nor lowered endogenous RPL22L1 protein levels **(Fig. 5e)**. Consistent with the underlying model, immunoprecipitation of ribosomes using HA-RPL7 recovered substantially more RPL22L1 from RPL22 KO cells than from control KO cells **(Supplementary Fig. 5d)**, indicating that RPL22L1 is incorporated into ribosomes to a greater extent when RPL22 is absent. These results suggest that RPL22L1 is recognized by BAG6 for proteasomal degradation when it fails to be incorporated into the ribosomal site shared with RPL22.

We next asked whether this mechanism operates in a physiologically relevant setting of reduced RPL22 expression. HCT116 colorectal carcinoma cells are microsatellite-instability-high and carry a heterozygous RPL22 frameshift mutation (p.K15Rfs*5), which introduces a premature stop codon and yields a truncated, non-functional protein^38,39^. Consistent with this genotype, HCT116 cells showed reduced RPL22 expression and correspondingly elevated levels of RPL22L1 relative to HEK293T cells **(Fig. 5f)**. Whereas BAG6 ablation in HEK293T cells increased RPL22L1 abundance, BAG6 loss had only a modest effect in HCT116 cells, where RPL22L1 levels were already elevated **(Fig. 5f)**, consistent with the idea that a larger fraction of RPL22L1 is incorporated into ribosomes and therefore protected from BAG6-mediated degradation.

Conversely, re-expression of HA-RPL22 in HCT116 cells reduced RPL22L1 abundance, whereas HA-RPL7 did not **(Fig. 5g)**. Because a decrease in steady-state levels could in principle reflect transcriptional or translational changes rather than enhanced turnover, we asked whether this reduction depends on proteasome activity. Treatment with the proteasome inhibitor bortezomib restored RPL22L1 levels in HA-RPL22-expressing cells, with TSPYL1 serving as a control for efficient proteasome inhibition **(Fig. 5h)**. Thus, restoration of WT RPL22 re-sensitizes RPL22L1 to BAG6-dependent proteasomal degradation in cancer cells that have naturally lost RPL22 expression.

Together, these findings define a post-translational mechanism that couples RPL22L1 abundance to ribosome occupancy **(Fig. 6)**. In cells expressing RPL22, the shared site on the 60S subunit is occupied by RPL22, leaving newly synthesized RPL22L1 unassembled. In this state, its bipartite hydrophobic degron is accessible to BAG6, which recruits RNF115 through its UBL domain to ubiquitinate RPL22L1 and direct it to the proteasome. When RPL22 is lost or mutated, a vacant binding site allows RPL22L1 incorporation into the 60S subunit, where it is protected from BAG6-RNF115-mediated degradation and enables its accumulation as a functional substitute for RPL22. These findings reveal a broader principle by which cells maintain ribosome proteostasis: unassembled ribosomal components are selectively eliminated, whereas productive assembly serves as a signal for stabilization and functional incorporation.

**Fig. 6.**
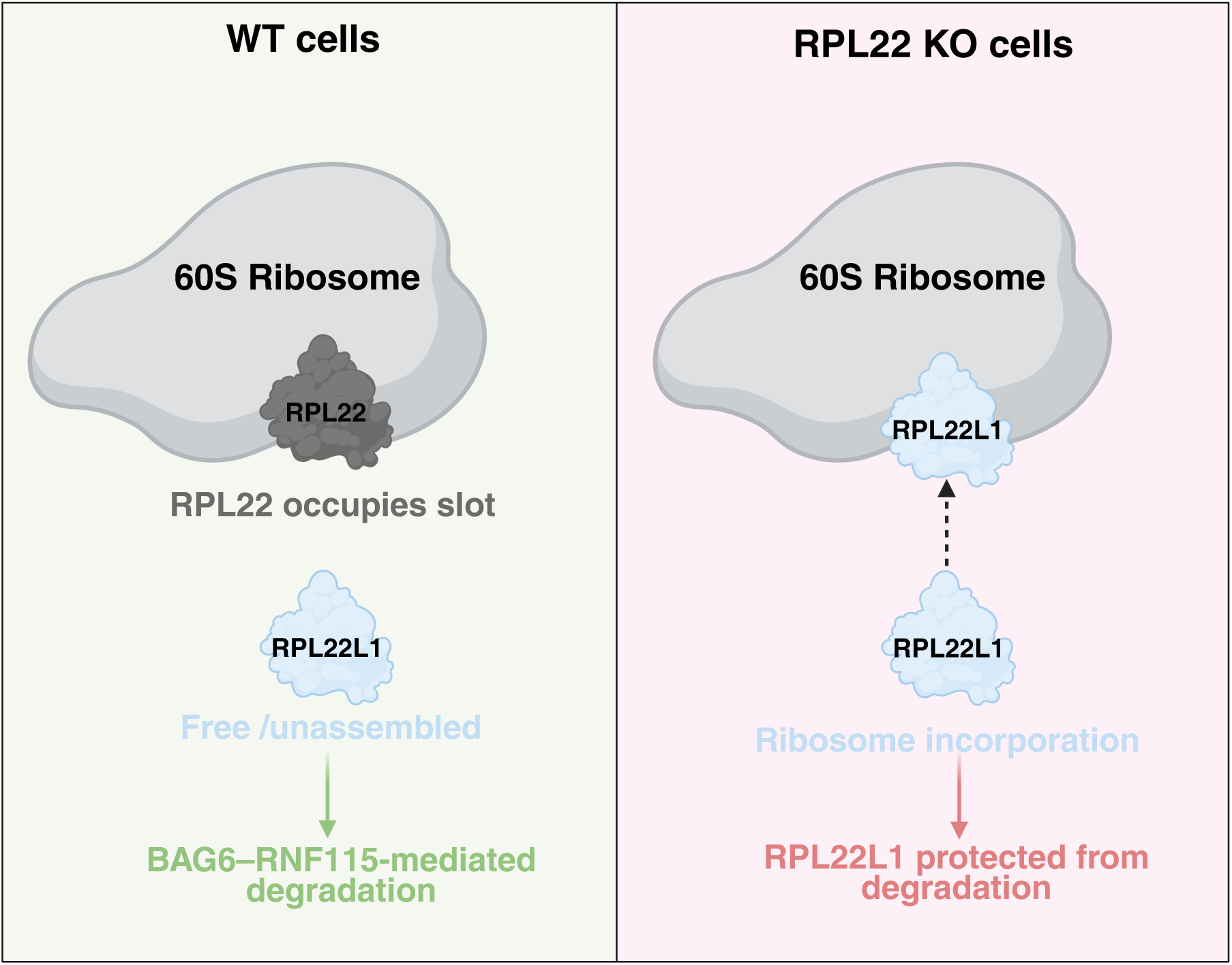
Model for unassembly-coupled degradation of RPL22L1 by the BAG6-RNF115 axis. Schematic of the proposed mechanism. In cells expressing RPL22, RPL22 occupies the shared binding site on the 60S ribosomal subunit, leaving newly synthesized RPL22L1 free and unassembled. In this state, the RPL22L1 degron is accessible to the BAG6 complex, which recruits RNF115 to ubiquitinate RPL22L1 and target it for proteasomal degradation. When RPL22 is absent or mutated, RPL22L1 is incorporated into the 60S subunit in its place; ribosome-associated RPL22L1 is protected from BAG6-RNF115-mediated degradation and accumulates to maintain cell functionality and proper ribosomal assembly.

## DISCUSSION

Ribosomal proteins are a major source of orphan quality control substrates, and multiple PQC pathways recognize distinct features exposed in unassembled or excess ribosomal proteins. The yeast E3 ligase Tom1 and its mammalian homolog HUWE1, for example, restrict accumulation of overproduced Rpl26 by recognizing residues that are buried in the mature ribosome^40^. In a parallel route, the E2-E3 hybrid enzyme UBE2O directly recognizes juxtaposed basic and hydrophobic patches that are normally shielded by nuclear import factors or within the assembled ribosome, targeting nascent ribosomal proteins that fail import or assembly^41^. Our findings add BAG6–RNF115 to this network and reveal a distinct mode of recognition, in which hydrophobic residues within a bipartite degron of RPL22L1 are recognized by BAG6 when the protein remains unassembled. Ribosome incorporation masks this degron and thereby protects RPL22L1 from degradation, coupling PQC to its assembly state.

Similar orphan quality control mechanisms operate on other multiprotein complexes, where a dedicated E3 ligase is directed to the unassembled subunit by an adaptor protein rather than binding it directly. The E3 ligase HERC1 targets unassembled PSMC5, a subunit of the proteasome base, via its cognate assembly chaperone PAAF1^42^. In an analogous manner, the ligase HERC2 has been shown to target orphan CCT subunits through the adaptor ZNRD2, which binds unassembled but not assembled chaperonin^42^. Our findings follow the same principle as BAG6 acts as the adaptor that directly recognizes an exposed degron and recruits RNF115 for further ubiquitination and degradation; this motif becomes inaccessible once RPL22L1 is assembled into the 60S subunit.

Our data indicates that RPL22L1 is selectively degraded by RNF115 rather than the closely related RNF126. Why RPL22L1 is preferentially handled by RNF115 remains unclear. Differential localization is unlikely to account for it, as both E3 ligases are present in the cytosol and the nucleus **(Fig. 3a)** and both are recruited through the same BAG6 UBL domain^22,26^. Instead, the distinction may reflect the nature of the RPL22L1 degron. RNF126 acts on strong hydrophobic BAG6 clients, such as the transmembrane domains of mislocalized membrane proteins^22^. RPL22L1 is a soluble, highly basic protein whose degron consists of short hydrophobic stretches distributed within two discrete regions rather than a continuous hydrophobic segment. These features may result in weaker BAG6 engagement, thereby requiring additional substrate recognition by RNF115. Consistent with this possibility, RNF115 can recognize some substrates independently of BAG6, such as tetherin, through a discrete region N-terminal to its RING domain^43^. Alternatively, RNF115 and RNF126 may differ in their E2 partners or in the ubiquitin chain architectures they generate, such that only RNF115 produces a degradation-competent ubiquitin signal on RPL22L1. Although the two E3 ligases share conserved N-terminal zinc-finger and C-terminal RING domains, they diverge in other regions **(Supplementary Fig. 2a-b)**, and domain-swap experiments between the RNF115 and RNF126 should help distinguish whether their substrate specificity arises from differential substrate engagement or from downstream ubiquitination activity.

Beyond its role in protein synthesis as a component of the ribosome, RPL22L1 has extraribosomal functions, acting as a regulator of pre-mRNA splicing^29^. In cells lacking RPL22, the accumulating RPL22L1 both occupies the vacant ribosomal binding site and promotes a splicing switch in MDM4 that favors the oncogenic full-length isoform, attenuating p53 activity and supporting tumor growth in microsatellite-unstable cancers^37^. Because splicing occurs in the nucleus, it is the nuclear pool of RPL22L1 that carries this activity. Our finding that BAG6 and RNF115 are both present in the nucleus **(Fig. 3a)**, together with the stabilization of nuclear RPL22L1 following BAG6 loss **(Fig. 3b-d)**, raises the possibility that BAG6-mediated degradation also regulates the nuclear, extraribosomal pool of RPL22L1. Thus, BAG6 surveillance may act across compartments to control not only the pool of RPL22L1 available for ribosome assembly but also the abundance of RPL22L1 available for extraribosomal functions such as splicing. Such dual-compartment surveillance could help explain the tight regulation of RPL22L1 by BAG6 even though the protein itself is fully functional.

While BAG6 regulates the abundance of one ribosomal protein paralog pair, whether similar paralog-selective quality control extends to other ribosomal protein paralogs remains an open question. Only a small number of human ribosomal proteins are encoded as paralog pairs, with each pair competing for the same site on the ribosome. Our findings suggest that sequence divergence between paralogs can itself encode a degron, providing a mechanism to selectively eliminate the unassembled species while sparing its ribosome-incorporated partner. Whether other paralog pairs are subject to comparable post-translational surveillance, and which E3 ligases mediate their turnover, remains to be determined. Notably, RPL22L1 was the sole ribosomal protein whose abundance increased upon BAG6 loss, suggesting that BAG6-mediated surveillance is highly selective and that other paralog pairs, if similarly regulated, may be controlled by distinct pathways.

Together, our findings identify BAG6 as a sensor of protein assembly state and suggest that unassembled-coupled degradation can serve as a means of matching protein abundance to functional demand.

## METHODS

### Cell culture

HEK293T cells (ATCC® CRL-3216™), U2OS (a gift from Shav-tal Y. lab, Bar-Ilan University), HCT116 (a gift from Cohen HY. lab, Bar-Ilan University) and HeLa cells (a gift from Kimchi A. lab, Weizmann Institute of Science) were grown in Dulbecco’s Modified Eagle’s Medium (DMEM) (Life Technologies) supplemented with 10% Fetal Bovine Serum (FBS) (Gibco) and 100 U ml^−1^ penicillin/streptomycin (Life Technologies or Gibco). All cells were maintained at 37 °C in a humidified incubator with 5% CO2 and were routinely tested for mycoplasma contamination.

### Transfection and lentivirus production

Lentivirus was generated through the transfection of HEK293T cells using PolyJet In Vitro DNA Transfection Reagent (SignaGen Laboratories). Cells seeded at approximately 80% confluency were transfected as recommended by the manufacturer with the lentiviral transfer vector plus four plasmids encoding Gag-Pol, Rev, Tat, and VSV-G. The media was changed 24 h post-transfection and lentiviral supernatants collected a further 24 hr later. Cell debris was removed by centrifugation (800 x g, 5 min) and virus was stored in single-use aliquots at −80°C. Transduction of target cells was achieved by adding the virus in the presence of 8 μg/ml hexadimethrine bromide (Polybrene).

### Inhibitors

Bortezomib (APExBIO, #A2614) and MLN7243 (Selleck Chemicals, #S8341) were used at a final concentration of 1 µM for 6 hr. Cycloheximide (CHX; APExBIO, #A8244) was used at a final concentration of 50 µg ml^−1^.

### Plasmids

The open reading frames (ORFs) encoding BAG6, RNF115, RNF126, RPL7, and RPL22 were obtained in the form of entry clones from the Ultimate ORF clone collection (Thermo Fisher Scientific). The indicated ORFs were cloned into a pHAGE N-terminal FLAG-HA destination vector by LR reaction (Gateway LR Clonase II Enzyme Mix; Thermo Fisher Scientific, 11791020) according to the manufacturer’s protocol. In addition, RPL7, RPL22L1 and RPL22 were also cloned into the pHAGE-GPS3.0^10^ destination vector by LR reaction. All constructs were verified by Sanger sequencing.

To generate the WT RPL22L1-HA vector (Fig. 2b), RPL22L1 was amplified by PCR to add HA tag to its C-terminus and subsequently cloned into the pHAGE Lenti Crimson vector (EF1α[promoter]-ORF-PGK[promoter]-Crimson)^44^ using HiFi DNA assembly. The BAG6 ΔUBL truncation variant, lacking the first 100 aa of the N-terminal ubiquitin-like (UBL) domain, was generated by PCR amplification followed by ligation into the same vector.

To generate the catalytically inactive RNF115 mutant, two zinc-coordinating cysteines within the RNF115 RING domain (C228 and C231) were substituted with alanine. Site-directed mutagenesis was performed directly on the pHAGE N-terminal FLAG–HA RNF115 expression construct using the QuikChange Lightning Kit (Agilent, #210518) according to the manufacturer’s protocol, with the primers indicated below. The mutations were confirmed by Sanger sequencing. The primer set used to generate the RNF115 C228A/C231A mutant for expression in mammalian cells was: RNF115 C228A/C231A Fw: ctcttcaactgtgtaatcttctttggctactggagcctctaaacccatatcaacttgttcc RNF115 C228A/C231A Rv: ggaacaagttgatatgggtttagaggctccagtagccaaagaagattacacagttgaagag

To map the region of RPL22L1 required for BAG6-dependent degradation, chimeric mutants were generated in which defined regions of RPL22L1 (residues 1-33, ‘N-terminal’; 39-71, ‘Middle’; and 99-122, ‘C-terminal’) were replaced with the corresponding sequences of RPL22, generating the (N)22, (M)22 and (C)22 chimeras, respectively. To test whether these regions are sufficient to confer degradation, reciprocal chimeras were generated in which the RPL22L1 N-terminal (residues 1-33) or Middle (residues 39-71) region was transplanted into the corresponding position of RPL22, generating RPL22(N)22L1 and RPL22(M)22L1. All constructs were designed on the basis of the canonical human sequences of RPL22L1 (UniProt Q6P5R6) and RPL22 (UniProt P35268). Full-length RPL22L1 and all chimeric ORFs were synthesized by GenScript.

To test the isolated degron regions, the RPL22L1 N-terminal peptide (residues 1-33), the Middle peptide (residues 39-71), and a combined N+M fragment (residues 1-71) were used as minimal reporter substrates. To identify the residues required for recognition, a degron mutant was generated in which the ten hydrophobic residues within the Middle degron (V42, V44, L51, V54, V55, I57, F60, I64, V66, and V67) were substituted with alanine. DNA encoding all peptide fragments was synthesized by Integrated DNA Technologies (IDT).

All fragments were cloned into the pHAGE-GPS3.0 lentiviral reporter vector, linearized with BstBI and XhoI, using NEBuilder HiFi DNA Assembly Master Mix (New England Biolabs, E2621L), such that each fragment was fused in frame to the C-terminus of GFP. All constructs were verified by Sanger sequencing.

CRISPR-Cas9-mediated gene disruption was performed using the lentiCRISPR v2 vector (Addgene, 52961). Oligonucleotides encoding the top and bottom strands of the sgRNAs were synthesized (Sigma-Aldrich), annealed, and cloned into the vector as previously described. Nucleotide sequences of the sgRNAs used in this study are as follows:

sg-AAVS1: GGGGCCACTAGGGACAGGAT
sg1-BAG6: GGAGGTGTTGGTGAAGACCT
sg2-BAG6: GAGGCTCCTCCACAGCGGTAC
sg3-BAG6: GCAAGATGATAAGAAGCTTC
sg-RNF115: GAAAGTGGCAGAAAAACCGGT
sg-RNF126: GCAAACTGTCCGTAGCCCTG
sg1-RPL22: GCAAGATCACCGTGACATCCG
sg2-RPL22: GCGCGGAGCCATACTAACCAC

### Generation of CRISPR-Cas9 KO cells

Lentivirus was generated by transfecting HEK293T cells with lentiCRISPR v2 encoding the relevant sgRNA, as described above. Forty-eight hours after transduction, cells were selected with 1 µg ml⁻¹ puromycin for 3 d. KO cells were validated by western blot analysis for loss of the target protein. BAG6, RNF115, RNF126, and RPL22 KO cells were generated by viral transduction, with sgRNA targeting the AAVS1 safe-harbor locus used to generate control (Ctrl) KO cells. To control for off-target effects, two independent KO lines were generated for BAG6 and RPL22 genes using distinct sgRNAs; in the case of BAG6, KO1 was generated by combining sg1 and sg2, while KO2 was generated using sg3.

### Global protein stability assay

The global protein stability (GPS) reporter assay was performed as previously described. In brief, each reporter was expressed from a bicistronic lentiviral vector (pCMV–DsRed–IRES–GFP–ORF) in which the ORF of interest is fused to the C-terminus of GFP and DsRed is co-expressed as an internal normalization control; GFP/DsRed ratios were measured by flow cytometry as a readout of reporter stability. The reporters used in this assay were: full-length RPL22L1, RPL22 and RPL7; the RPL22L1-based chimeric mutants (N)22, (M)22 and (C)22, in which the N-terminal (residues 1-33), Middle (residues 39-71) or C-terminal (residues 99-122) region of RPL22L1 was replaced with the corresponding region of RPL22; the reciprocal RPL22-based chimeric mutants RPL22 (N)22L1 and RPL22 (M)22L1, in which the corresponding N-terminal or Middle region of RPL22 was replaced with that of RPL22L1; Isolated GFP-fused N-terminal (residues 1-33), Middle (residues 39-71) and N+M (residues 1-71) peptides of RPL22L1; a degron-mutant reporter in which the hydrophobic residues within the RPL22L1 Middle degron were substituted with alanine; and the nuclear-restricted NLS-GPS-RPL22L1 reporter. Construct details are described in the plasmids and cloning section.

Lentiviral particles encoding the GPS reporters were introduced into WT or KO cells, followed by blasticidin selection (20 μg ml^−1^) for 3 d. To assess the effect of cDNA rescue, stable GPS reporter cells were subsequently transduced with lentiviruses encoding HA-BAG6 (WT or ΔUBL), HA-RNF115 (WT or C228A/C231A mutant), HA-RPL7, or HA-RPL22, followed by puromycin selection (1 μg ml^−1^) for 3 d. Flow cytometry was performed on a CytoFLEX (Beckman Coulter) to record GFP and DsRed fluorescence, and the GFP/DsRed ratio, reporting on substrate stability, was analyzed using FlowJo v.10.9. Where indicated, cells were treated for 6 hr with 1 µM bortezomib or 1 µM MLN7243 prior to analysis.

### Immunoblotting

Cells were lysed in ice-cold lysis buffer (10 mM NaPO4, 100 mM NaCl, 5 mM EDTA (pH 8), 1% Triton X-100, 0.5% deoxycholic acid sodium salt and 0.1% SDS) supplemented with Halt Protease and Phosphatase Inhibitor Cocktail (Thermo Fisher Scientific, 78442) for 25 min at 4 °C. Lysates were clarified by centrifugation (20,000g, 15 min, 4 °C). Protein concentration was determined by Bradford assay (Bio-Rad, 500-0006). Proteins were resolved by SDS-PAGE (Mini-PROTEAN TGX Precast Protein Gels; Bio-Rad) and transferred to a nitrocellulose membrane, which was blocked in 10% non-fat dry milk in PBS + 0.1% Tween 20 (PBS-T). Membranes were incubated with primary antibody overnight at 4 °C, washed three times with PBS-T, and then incubated with HRP-conjugated secondary antibodies, goat anti-rabbit IgG (H+L) HRP (1:20,000; Jackson ImmunoResearch, 111-035-144) or goat anti-mouse IgG (H+L) (1:20,000; Jackson ImmunoResearch, 115-035-003) for 1 h at room temperature. Reactive bands were visualized using SuperSignal West Femto Chemiluminescent Substrate (Pierce, 34095) on an Amersham Imager 680 (Cytiva) using ImageQuant TL software.

The following primary antibodies were used: mouse anti-vinculin (1:1,000; Sigma-Aldrich, V9264), rabbit anti-HA tag (1:1,000; Cell Signaling Technology (CST), 3724), rabbit anti-BAG6 (1:1,000; Cell Signaling Technology (CST), 8523), rabbit anti-RNF115 (1:1,000; Abcam, ab187642), rabbit anti-RNF126 (1:1,000; Proteintech, 66647-1-Ig), rabbit anti-RPL22 (1:1,000; Cell Signaling Technology (CST), 54055), rabbit anti-RPL22L1 (1:1,000; Cell Signaling Technology (CST), 40259), rabbit anti-RPL7 (1:1,000; Bethyl Laboratories, A300-741A), rabbit anti-RPS3 (1:1,000; Bethyl Laboratories, A303-841A), rabbit anti-TSPYL1 (1:5,000; Bethyl Laboratories, A304-852A), rabbit anti-GAPDH (1:1,000; Cell Signaling Technology (CST), 2118), rabbit anti-β-tubulin (1:1,000; Cell Signaling Technology (CST), 2146), rabbit anti-Histone H3 (1:1,000; Cell Signaling Technology (CST), 9715), rabbit anti-RPS6 (1:1,000; Bethyl Laboratories, A300-556A), rabbit anti-GFP (1:1000, Abcam, ab290), mouse anti-B23/nucleophosmin (1:5000; Abcam ab86712).

### Immunoprecipitation

HEK293T cells stably expressing HA-BAG6 (WT or ΔUBL), HA-RNF115, HA-RNF126, RPL22L1-HA or HA-RPL7 were generated by lentiviral transduction. For IP experiments, cells on 10-cm plates were pre-treated with 1 μM bortezomib for 6 hr. Cells were lysed in NP-40 lysis buffer (0.5% NP-40, 50 mM Tris-HCl pH 7.4, 150 mM NaCl) supplemented with protease and phosphatase inhibitors. Cleared lysates were incubated with anti-HA Magnetic Beads (Sigma-Aldrich, SAE0197), and immunoprecipitates were washed three times in wash buffer (0.5% NP-40, 50 mM Tris-HCl pH 7.4, 300 mM NaCl) supplemented with protease and phosphatase inhibitors. For the HA-RPL7 co-immunoprecipitation (Supplementary Fig. 5c), both the lysis and wash buffers were additionally supplemented with 5 mM MgCl₂ to preserve ribosomal interactions. Bound proteins were eluted by incubation with HA peptide (Sigma-Aldrich, I2149) in elution buffer (50 mM Tris-HCl pH 7.4, 150 mM NaCl) for 30 min at 37 °C with rotation. The cleared supernatant was collected, and eluted proteins were denatured in SDS-PAGE sample buffer (95 °C, 10 min) prior to immunoblotting.

For immunoprecipitation of GFP-RPL22 in Supplementary Fig. 1e, HEK293T cells stably expressing GFP-RPL22 were pre-treated with 1 μM bortezomib for 6 h, followed by lysis in ice-cold NP-40 lysis buffer (0.5% NP-40, 50 mM Tris-HCl pH 7.4, 150 mM NaCl) supplemented with protease and phosphatase inhibitors and 5mM MgCl₂ to preserve ribosomal interactions. After 30 min on ice, lysates were clarified by centrifugation (20,000 × g, 10 min, 4°C) followed by immunoprecipitation using GFP-Trap MA magnetic agarose beads (Chromotek, gtma), which were added to the supernatants and incubated with rotation for 2 hr at 4°C. The beads were then washed three times with wash buffer (0.5% NP-40, 50 mM Tris-HCl pH 7.4, 300 mM NaCl) before bound proteins were eluted upon incubation with SDS-PAGE sample buffer (95°C, 10 min). Proteins were subsequently resolved by SDS-PAGE as explained before.

### Nuclear-cytoplasmic fractionation

HEK293T cells (3 × 10⁶) were harvested by trypsinization, washed once in ice-cold PBS, and pelleted at 900 × g for 5 min. The cell pellet was lysed in digitonin lysis buffer (500 µg/ml digitonin, 20 mM HEPES pH 7.3, 110 mM potassium acetate, 2 mM magnesium acetate, 5 mM sodium acetate, 2 mM EGTA-NaOH pH 7.3) supplemented with protease and phosphatase inhibitors for 1 min on ice. Cells were immediately centrifuged at 900 × g for 5 min, and the cleared supernatant, containing the cytosolic fraction, was retained and re-centrifuged at 900 × g for 5 min to remove residual nuclei. The nuclei-containing pellet was washed once in wash buffer (20 mM HEPES pH 7.3, 110 mM potassium acetate, 2 mM magnesium acetate, 5 mM sodium acetate, 2 mM EGTA-NaOH pH 7.3) supplemented with protease inhibitors (900 × g, 5 min). The washed nuclear pellet was lysed in urea buffer (10 mM Tris-HCl pH 7.4, 150 mM NaCl, 1 mM EDTA pH 8.0, 1 mM EGTA, 0.5% Triton X-100, 7 M urea) and sonicated for a total of 60 s (30 s on/30 s off, 40% amplitude). Lysates were incubated on ice for 30 min and then centrifuged at 17,000 × g for 10 min to yield the nuclear fraction. Protein concentrations were determined, and equal amounts of each fraction were resolved by SDS-PAGE. Fractionation efficiency was confirmed by immunoblotting for β-tubulin (cytosolic marker) and B23 or histone H3 (nuclear markers).

### Cycloheximide chase assay

After treatment with 50 μg ml^−1^ CHX (APExBIO, A8244), cells were harvested at the indicated time points (0, 12 and 24 hr) and subjected to immunoblotting as described above. TSPYL1, a known short-lived CRL2 substrate, served as an internal positive control for efficient translation inhibition, and vinculin or GAPDH served as a loading control.

### Quantitative RT-PCR

Cells were harvested, and total RNA was isolated using a Direct-zol RNA Microprep Kit (Zymo Research, R2061), followed by cDNA synthesis with a qScript cDNA Synthesis Kit (Quantabio, 95047-025). Quantitative PCR was performed using Power SYBR Green PCR Master Mix (Applied Biosystems, AB-4367659) on a CFX Connect Real-Time PCR Detection System (Bio-Rad). Relative expression of RPL22L1 was calculated using the ΔΔCt method and normalized to β-actin. Data represent three independent experiments shown as mean ± s.e. of n = 3 technical replicates, and P values were determined by a two-tailed unpaired t-test. The primers used were as follows:

RPL22L1 Fw: AGACAGGAAGCCCAAGAGGT
RPL22L1 Rv: TCCCGAGATTTCCAGTTTTG
β-actin Fw: CACCTTCTACAATGAGCTGCGTGTG
β-actin Rv: ATAGCACAGCCTGGATAGCAACGTAC

### Confocal microscopy

HEK293T cells stably expressing GFP-RPL22L1 (WT, (N)22, (M)22 or (C)22 chimera) within the GPS bicistronic vector (co-expressing DsRed) were grown on coverslips, fixed for 15 min with 4% formaldehyde, and permeabilized and blocked with 3% BSA and 0.5% Triton X-100 in PBS. To visualize the nucleolus, coverslips were incubated overnight at 4 °C with mouse anti-B23/nucleophosmin antibody (1:200; Abcam ab86712), followed by goat anti-mouse IgG Alexa Fluor 647 (1:500; Abcam, ab175473) for 1 hr at room temperature. Nuclei were counterstained with DAPI (Sigma-Aldrich, D9542) for 1 min. Coverslips were mounted onto slides (Sigma-Aldrich, F6182) and imaged with a Leica Stellaris 5 confocal microscope using LAS X software and a ×63 oil/1.4 NA objective. GFP-RPL22L1 was visualized in the green channel and DsRed (cytoplasmic control) in the yellow channel.

### Image quantification of cytoplasmic and nuclear GFP-RPL22L1 fluorescence

GFP-RPL22L1 fluorescence was quantified at the field level in Fiji/ImageJ (version 2.14.0/1.54h) using a custom macro. For each two-channel image (DsRed, GFP), both channels were background-subtracted (rolling-ball radius, 50 px). Because DsRed fills the cytoplasm but is largely excluded from the nucleus, the DsRed channel was used to define subcellular regions by intensity thresholding: a high DsRed threshold selected the bright cytoplasmic region (excluding the dim nuclear centers), and a lower DsRed threshold selected the entire cell footprint (including nuclei). The nuclear region was defined as the difference between the whole-cell and cytoplasmic selections (whole-cell XOR cytoplasm). Both DsRed thresholds were set manually on the first image and applied identically to all images across every condition within an experiment, and identical acquisition settings were used throughout. Mean GFP intensity was then measured within the cytoplasmic and nuclear regions of each field. Cytoplasmic and nuclear signals were analyzed separately; within each compartment, per-field GFP intensity was normalized to the mean intensity of the Ctrl KO condition, such that the control mean was set to 1, and BAG6 KO values were expressed relative to control. Data are presented as relative GFP–RPL22L1 intensity (mean ± s.d.), with each point representing one imaging field (n = 4 per condition). Statistical significance was assessed by a two-tailed unpaired t-test

### Proteomics

#### Cell lysis and protein digestion

HEK293T WT cells and two independent BAG6 KO lines were seeded in 10-cm dishes and grown for 48 h before harvest. After aspiration of media, cells were rinsed (3×) with ice-cold PBS and lysed by adding 600 µl of ice-cold lysis buffer (RIPA buffer supplemented with 1× Protease and Phosphatase Inhibitor Cocktail, 10 mM sodium pyrophosphate, 10 mM β-glycerophosphate, 0.2 mM TCEP and 100 µM PR-619). Lysates were scraped, homogenized, incubated on ice for 5 min, sonicated and clarified by centrifugation at 20,000g (5 min, 4 °C). Protein concentrations were determined by Bradford assay. Protein extracts (100 µg) underwent disulfide bond reduction with 5 mM TCEP (10 min) and alkylation with 20 mM iodoacetamide (15 min). Methanol–chloroform precipitation was performed before overnight trypsin digestion at a 100:1 protein-to-protease ratio in 100 mM EPPS (pH 8.5) containing 0.1% RapiGest.

#### TMT labeling

TMT reagent was added with acetonitrile to a final concentration of approximately 30% (v/v), and after 1 h at room temperature, the reaction was quenched with hydroxylamine (final 0.5% v/v, 15 min). Labeling efficiency was confirmed, and labeled samples were pooled at a 1:1 ratio, vacuum centrifuged to near dryness, and subjected to C18 solid-phase extraction (SPE).

#### Off-line basic-pH reversed-phase fractionation

The pooled TMT-labeled sample was resuspended in 100 μl of 10 mM NH_4_HCO_3_ (pH 8.0) and fractionated by basic-pH reversed-phase HPLC (Agilent LC1260) through an Aeris PEPTIDE XB-C18 column (Phenomenex) into 96 fractions over a 90-min gradient, which were then concatenated into 24 non-contiguous fractions. Each fraction was desalted by StageTip, vacuum centrifuged, and reconstituted in 5% acetonitrile, 1% formic acid for LC–MS/MS analysis.

#### MS analysis

MS data were collected on an Orbitrap Eclipse Tribrid mass spectrometer coupled to an UltiMate 3000 RSLCnano LC pump. Peptides were separated on a 100-µm-inner-diameter microcapillary column packed in-house with approximately 40 cm of HALO Peptide ES-C18 resin (2.7 µm, 160 Å; Advanced Materials Technology), using a 105-min gradient at approximately 350 nl min^−1^. Each analysis used the Multi-Notch MS^3^-based TMT method^45^ to reduce ion interference compared to MS^2^ quantification^46^, combined with the FAIMS Pro Interface (using previously optimized 3 CV parameters for TMT-multiplexed samples^47^) and the newly implemented Real-Time Search analysis software^48–50^. The scan sequence began with an MS^1^ spectrum (Orbitrap analysis; resolution 60,000 at 200 Th; mass range 375−1500 m/z; automatic gain control (AGC) target 4×10^5^; maximum injection time 50 ms). Precursors for MS^2^ analysis were selected using a 1.25 sec/CV cycle type. MS^2^ analysis consisted of collision-induced dissociation (quadrupole ion trap analysis; Rapid scan rate; AGC 1.0×10^4^; isolation window 0.5 Th; normalized collision energy (NCE) 35; maximum injection time 35 ms). Monoisotopic peak assignment was used, and previously interrogated precursors were excluded using a dynamic window (180 s ±10 ppm). Following the acquisition of each MS^2^ spectrum, a synchronous-precursor-selection (SPS) API-MS^3^ scan was collected on the top 10 most intense ions b or y-ions matched by the online search algorithm in the associated MS^2^ spectrum^48–50^. MS^3^ precursors were fragmented by high-energy collision-induced dissociation (HCD) and analyzed using the Orbitrap (NCE 65; AGC 2.5×10^5^; maximum injection time 200 ms, resolution was 50,000 at 200 Th). The closeout was set to two peptides per protein per fraction, so MS^3^s were no longer collected for proteins with two peptide-spectrum matches (PSMs) that passed the quality filters.

#### Data analysis

Mass spectra were converted to mzXML^51^ and processed using the Comet search engine^52^ (2020.01 rev. 4) against the Human Reference Proteome (2024-05 release) with contaminants and reverse decoy sequences appended. Searches were performed with a 50-ppm precursor ion tolerance. TMT tags on lysine residues and peptide N-termini (+229.1629 Da) and carbamidomethylation of cysteine residues (+57.021 Da) were set as static modifications, and oxidation of methionine (+15.995 Da) was set as a variable modification. Peptide-spectrum matches (PSMs) for each run were adjusted to a 1% false discovery rate (FDR)^53^. PSM filtering was performed using a linear discriminant analysis employing a target-decoy strategy, while considering the following parameters: Comet Log Expect, ΔCn, charge state, missed cleavages, fraction of ions matched, precursor mass accuracy, and peptide length. Filtered PSMs were then further collapsed using the Picked FDR method^54^ to achieve a target FDR of 1% for each plex. Moreover, protein assembly was guided by principles of parsimony to produce the smallest set of proteins necessary to account for all observed peptides. TMT reporter ion intensities were extracted (integration tolerance of 0.003 Da) and corrected for isotopic impurities, and proteins were quantified by summing signal-to-noise across PSMs. Protein quantification values were exported for analysis in Microsoft Excel, Perseus, GraphPad Prism 10, and R 4.3.2. Differentially abundant proteins were identified between control parental (WT) HEK293T cells and the combined BAG6 KO lines using a two-sample Welch’s t-test, and P values were corrected for multiple testing by the Benjamini-Hochberg procedure (5% FDR). Proteins with a Benjamini–Hochberg-adjusted FDR < 5% and an absolute fold change > 2 (|log₂ fold change| > 1) were considered significantly altered, yielding 31 increased and 20 decreased proteins out of 9,940 quantified.

### Statistics and reproducibility

All statistical analyses were performed using GraphPad Prism 10. Statistical parameters, including the definition and exact values of n, distribution, and deviation, are reported in the figures and corresponding legends. Statistical significance was assessed using a two-tailed unpaired t-test unless otherwise indicated, with P > 0.05 considered not significant (NS) and P < 0.05 considered significant. Unless otherwise specified, all experiments were independently repeated at least three times with similar results, and representative results are shown throughout all figures.

## Illustrations

The graphical abstract (Fig. 6) and illustrations in Fig. 1f, Fig. 4a,c,f and Supplementary Fig. 4b were created with BioRender.

## Data and code availability

– All unique identifiers and web links for publicly available datasets are presented within the main text or the supplementary materials. Mass spectrometry data associated with Supplementary Table 1 have been deposited to the MassIVE repository and are publicly available as of the date of publication, with the dataset identifier MSV000102686.
– This paper does not report any original code.
– Any additional information required to reanalyze the data reported in this paper is available from the lead contact upon request.

## Acknowledgements

This work was supported by the Israel Science Foundation (ISF grants 2380/21 and 3096/21) (I.K.); the European Research Council (ERC) under the European Union’s Horizon 2020 research and innovation programme (ERC-2020-STG; grant agreement No. 947709) (A.R. and I.K.); the Pew Charitable Trusts (A.O), and the National Institutes of Health R35GM156454 (A.O.). C.Z. is supported by a Gerstner Sloan Kettering Beatrice P. K. Palestin Fellowship. This manuscript is the result of funding in part by the National Institutes of Health (NIH). It is subject to the NIH Public Access Policy. Through acceptance of this federal funding, NIH has been given a right to make this manuscript publicly available in PubMed Central upon the Official Date of Publication, as defined by NIH.

## Author Contributions

Conceptualization, A.R. and I.K.; Supervision, I.K.; Methodology, A.R. and I.K.; Investigation, A.R.; Formal analysis, A.R.; Proteomics, C.Z. and A.O.; Writing-original draft, A.R. and I.K.; Writing-review & editing, all authors.

## Competing Interest Statement

The authors declare that they do not have any conflicts of interest with the content of this article.

## Notes

### Competing Interest Statement

The authors have declared no competing interest.

