## Supplementary Figures for "BAG6-RNF115 Couples Protein Quality Control with Ribosome Assembly"

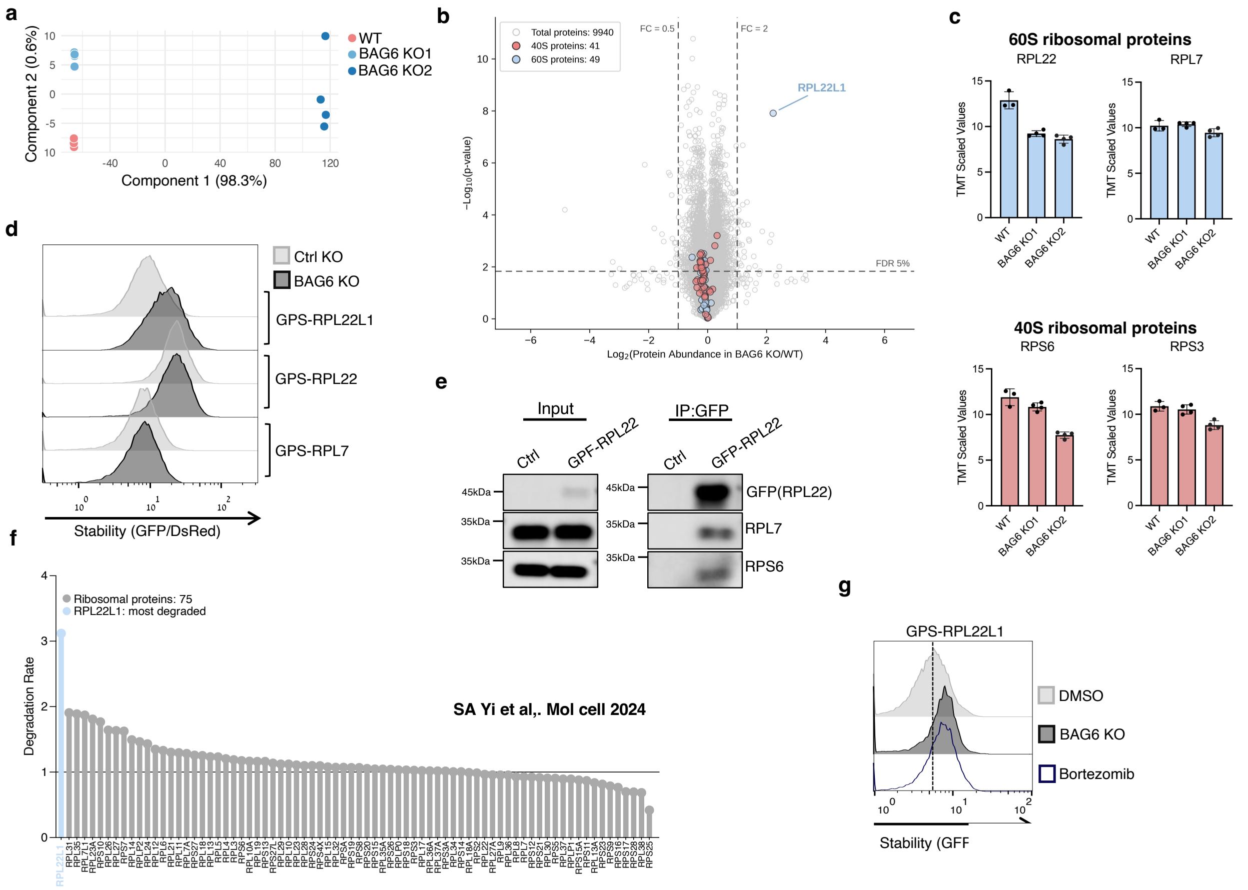


Supplementary Fig. 1 **TMT-based quantitative proteomics identifies RPL22L1 as the sole ribosomal protein targeted by BAG6.**

**a,** Principal component analysis (PCA) showing separate clustering of the control parental line (WT) and the two independent BAG6 KO HEK293T cell lines. Component 1, 98.3%; component 2, 0.6% of variance.

**b,** Volcano plot of the TMT-LC-MS proteome comparison (as in Fig. 1a). The 90 identified ribosomal proteins are highlighted according to ribosomal subunit, with 40S protein in salmon (n = 41) and 60S protein in light blue (n = 49), among 9,940 quantified proteins. Dashed lines indicate the fold-change (|FC| > 2) and FDR 5% thresholds. RPL22L1 is indicated as the only ribosomal protein exceeding both thresholds.

**c,** TMT relative abundance of representative 60S (RPL22, RPL7) and 40S (RPS6, RPS3) ribosomal proteins in WT and the two independent BAG6 KO lines. Data are mean ± s.e.; *n* = 3 (WT) and *n* = 4 (each BAG6 KO) biological replicates.

**d,** GPS stability analysis of GPS reporters fused to RPL22L1, RPL22 or RPL7 in Ctrl KO and BAG6 KO1 cells. RPL22L1 is the only reporter substantially stabilized upon BAG6 loss.

**e,** GFP immunoprecipitation of cells stably expressing GFP-RPL22 or empty vector (Ctrl) followed by immunoblotting for the endogenous ribosomal proteins RPL7 and RPS6. Input and immunoprecipitated samples (IP:GFP) are shown.

**f,** Re-analysis of the degradomics dataset of Yi et al., in which protein abundance was measured by quantitative proteomics at multiple time points following metabolic label washout (0, 5, 10 and 15 hr). For each ribosomal protein, the degradation rate was calculated as the ratio of abundance at 15 hr to 0 hr. Ribosomal proteins (*n* = 75) are plotted in descending order of degradation rate; RPL22L1 is highlighted in light blue as the most rapidly degraded ribosomal protein in the dataset. **g,** Flow cytometry analysis of the GPS-RPL22L1 reporter in cells treated with DMSO or 1 μM bortezomib for 6 hr, compared with BAG6 KO1 cells.

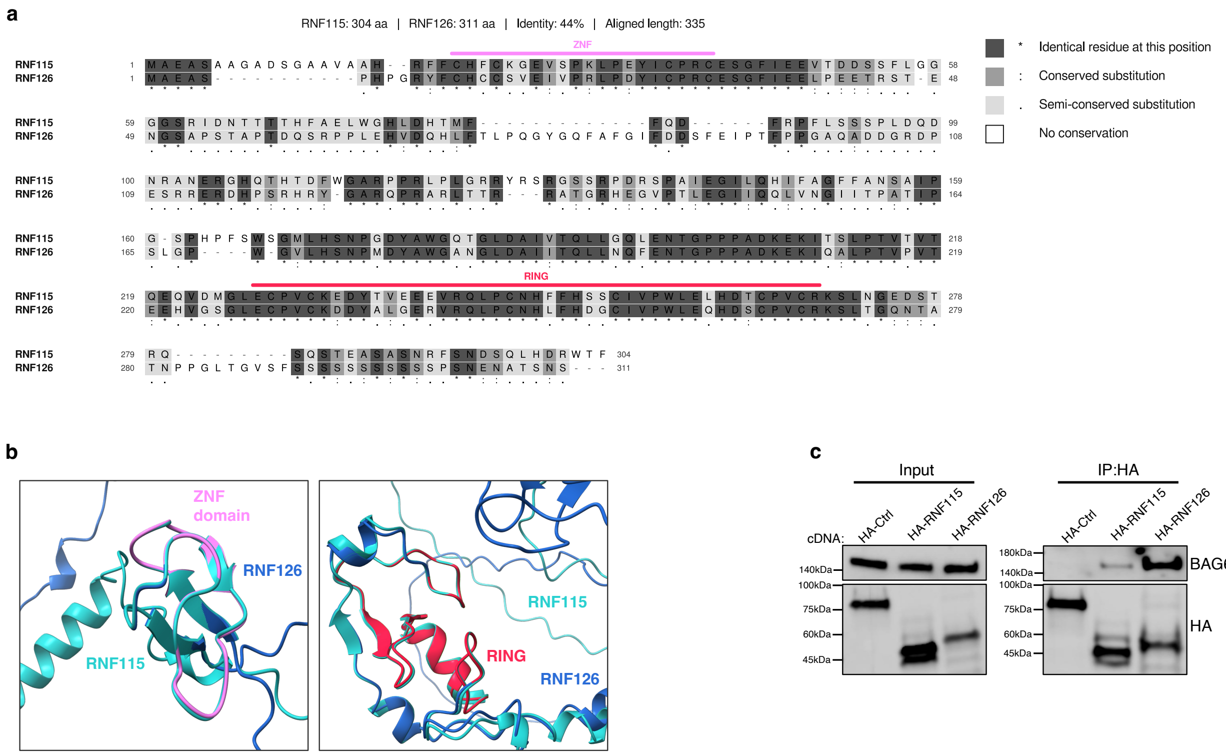


Supplementary Fig. 2 **RNF115 and RNF126 share conserved ZNF and RING domains and associate with BAG6.**

**a,** Pairwise sequence alignment of human RNF115 (UniProt Q9Y4L5, 304 aa) and RNF126 (UniProt Q9BV68, 311 aa), generated using EMBOSS Needle (44% identity over 335 aligned positions). Identical residues are shaded dark grey and marked with an asterisk (*); conserved substitutions, medium grey and colon (:); semi-conserved substitutions, light grey and period (.). Bars indicate the N-terminal C2/C2 zinc-finger (ZNF, pink) and C-terminal RING (red) domains.

**b,** Structural superposition of AlphaFold 3-predicted models of RNF115 (cyan) and RNF126 (blue), showing the ZNF domain (left, pink) and the RING domain (right, red). Visualized in ChimeraX.

**c,** HEK293T cells expressing either HA-tagged RNF115 or HA-tagged RNF126 were subjected to HA immunoprecipitation (IP:HA). Association with endogenous BAG6 was assessed by immunoblotting of immunoprecipitates and total cell extracts (Input). HA-Ctrl: HA-FEM1C cDNA.

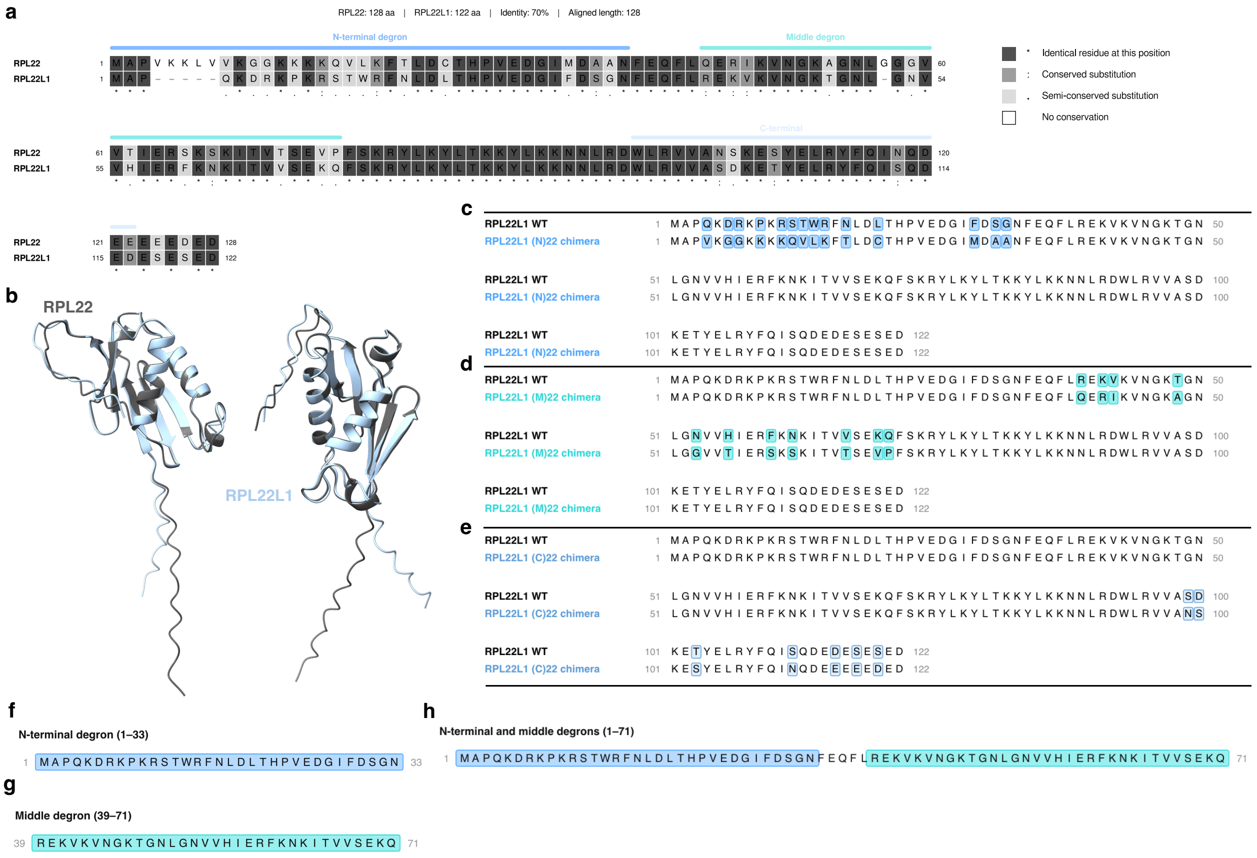


Supplementary Fig. 3 **Protein sequence comparison of RPL22L1 and RPL22 and design of degron-mapping constructs.**

**a,** Pairwise sequence alignment of human RPL22 (UniProt P35268, 128 aa) and RPL22L1 (UniProt Q6P5R6, 122 aa), generated using EMBOSS Needle (70% identity over 128 aligned positions). Identical residues are shaded dark grey and marked with an asterisk (*); conserved substitutions, medium grey and colon (:); semi-conserved substitutions, light grey and period (.). Bars indicate the N-terminal degron (blue) and middle degron (teal).

**b,** Structural superposition of AlphaFold 3-predicted models of RPL22 (dark grey) and RPL22L1 (light blue), shown in two orientations. Visualized in ChimeraX.

**c-e,** Chimeras are shown as alignments to wild-type RPL22L1 (122 aa) with residues differing from WT boxed; the (N)22 (blue), (M)22 (teal), and (C)22 (light blue) constructs carry swaps in the N-terminal (1-33), middle (39-71), and C-terminal (99-122) regions, respectively.

**f-h,** Degron constructs show the N-terminal (1-33), middle (39-71), and combined N-terminal + middle (1-71) sequences, shaded by region. Residue numbering follows WT RPL22L1.

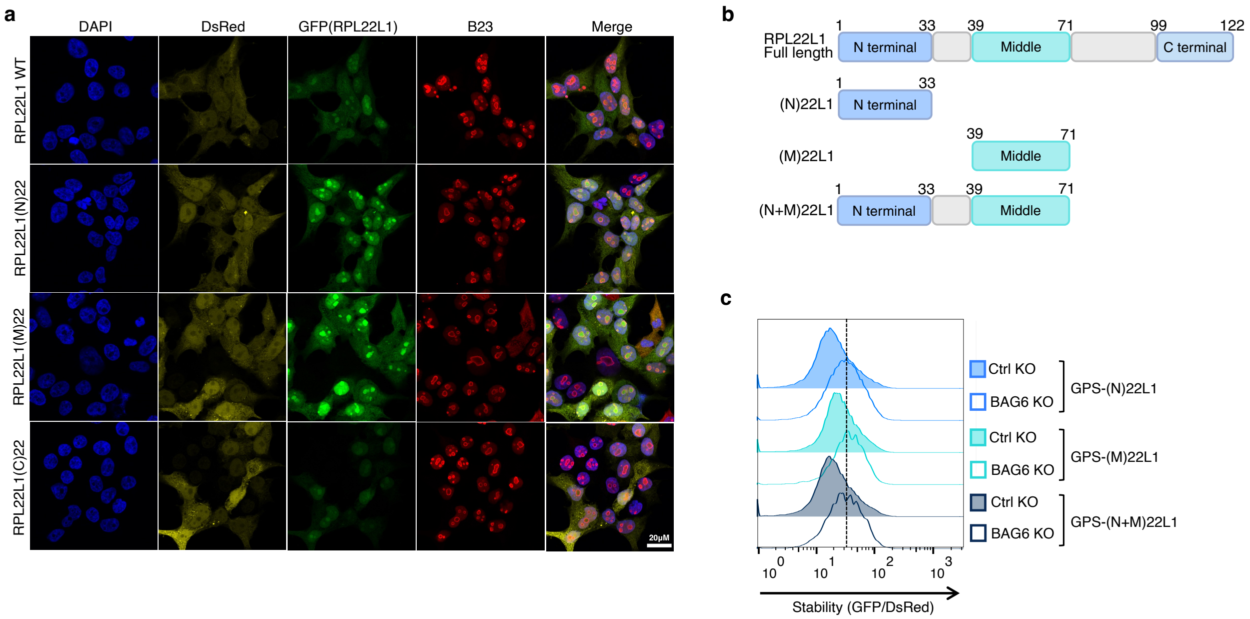


Supplementary Fig. 4 **Subcellular localization of GFP-RPL22L1 chimeras and autonomous degron activity of isolated regions.**

**a,** Representative confocal microscopy images of cells stably expressing GPS reporters with GFP-fused RPL22L1 WT, (N)22, (M)22 or (C)22. DAPI (blue), DsRed (yellow), GFP-RPL22L1 (green) and the nucleolar marker B23 (red) are shown with the merged image. Scale bar, 20 µm.

**b,** Schematic illustration of the isolated N-terminal region ((N)22L1, aa 1-33), middle region ((M)22L1, aa 39-71), and the combined N+M region of RPL22L1 ((N+M)22L1, aa 1-71), each fused to GFP within the GPS reporter.

**c,** Stability analysis of the GPS-(N)22L1, GPS-(M)22L1 and GPS-(N+M)22L1 reporters in Ctrl KO and BAG6 KO cells.

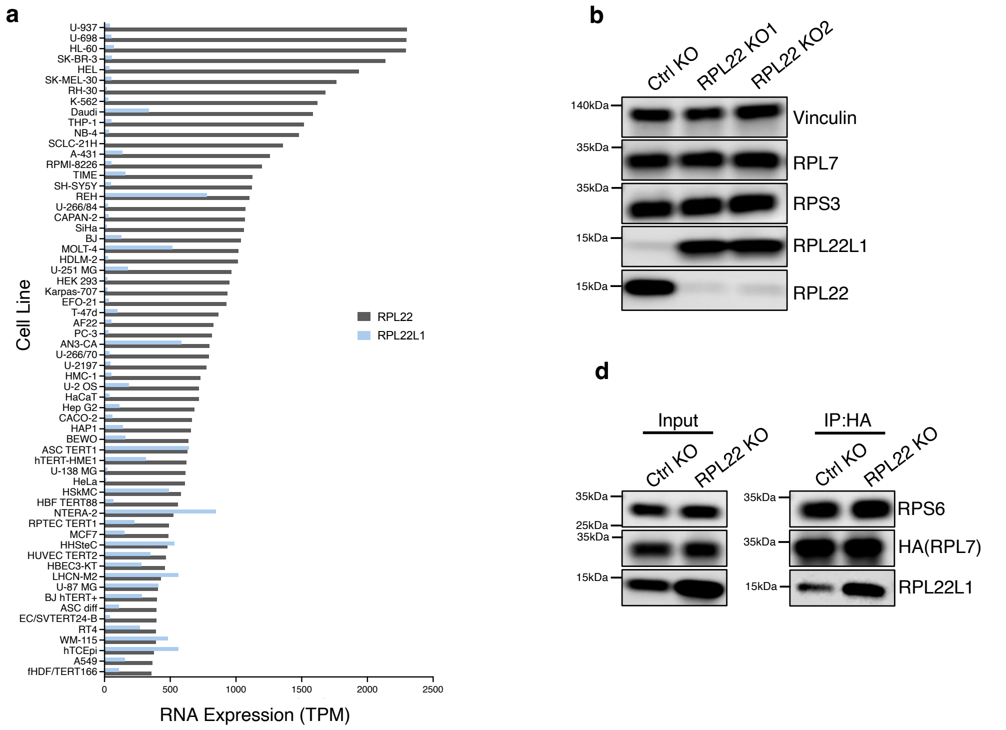


Supplementary Fig. 5 **RPL22 is the predominant paralog and limits RPL22L1 abundance by competing for ribosome incorporation.**

**a,** RNA expression of RPL22 (dark grey) and RPL22L1 (light blue) across human cell lines, in transcripts per million (TPM). Data obtained from the Human Protein Atlas.

**b,** Immunoblot analysis of the ribosomal proteins RPL7, RPS3, RPL22 and RPL22L1 in Ctrl KO and two independent RPL22 KO lines. Vinculin served as a loading control.

**c,** Quantitative RT-PCR of RPL22L1 mRNA in Ctrl KO and RPL22 KO cells. Data are mean ± s.e. of $n$ = 6 technical replicates, representative of two independent experiments. $P$ value determined by two-tailed unpaired $t$-test.

**d,** HA immunoprecipitation from Ctrl KO and RPL22 KO cells expressing HA-RPL7, followed by immunoblotting for RPS6, HA(RPL7) and RPL22L1 in immunoprecipitates and total cell extracts (Input).
